# Enhanced yield and subtype identity of hPSC-derived midbrain dopamine neuron by modulation of WNT and FGF18 signaling

**DOI:** 10.1101/2025.01.06.631400

**Authors:** Tae Wan Kim, Jinghua Piao, Vittoria D Bocchi, So Yeon Koo, Se Joon Choi, Fayzan Chaudhry, Donghe Yang, Hyein S Cho, Emiliano Hergenreder, Lucia Ruiz Perera, Subhashini Joshi, Zaki Abou Mrad, Nidia Claros, Shkurte Ademi Donohue, Anika K. Frank, Ryan Walsh, Eugene V. Mosharov, Doron Betel, Viviane Tabar, Lorenz Studer

## Abstract

While clinical trials are ongoing using human pluripotent stem cell-derived midbrain dopamine (mDA) neuron precursor grafts in Parkinson’s disease (PD), current protocols to derive mDA neurons remain suboptimal. In particular, the yield of TH+ mDA neurons after *in vivo* grafting and the expression of some mDA neuron and subtype-specific markers can be further improved. For example, characterization of mDA grafts by single cell transcriptomics has yielded only a small proportion of mDA neurons and a considerable fraction of contaminating cell populations.

Here we present an optimized mDA neuron differentiation strategy that builds on our clinical grade (“Boost”) protocol but includes the addition of FGF18 and IWP2 treatment (“Boost+”) at the mDA neurogenesis stage. We demonstrate that Boost+ mDA neurons show higher expression of EN1, PITX3 and ALDH1A1. Improvements in both mDA neurons yield and transcriptional similarity to primary mDA neurons is observed both in vitro and in grafts. Furthermore, grafts are enriched in authentic A9 mDA neurons by single nucSeq. Functional studies *in vitro* demonstrate increased dopamine production and release and improved electrophysiological properties. *In vivo* analyses show increased percentages of TH+ mDA neurons resulting in efficient rescue of amphetamine induced rotation behavior in the 6-OHDA rat model and rescue of some motor deficits in non-drug induced assays, including the ladder rung assay that is not improved by Boost mDA neurons. The Boost+ conditions present an optimized protocol with advantages for disease modeling and mDA neuron grafting paradigms.

## Introduction

Parkinson’s disease (PD) is the second most common neurodegenerative disease. It is characterized by the progressive degeneration of midbrain dopamine (mDA) neurons resulting in characteristic motor symptoms in PD patients, such as tremor, rigor, and bradykinesia(1, 2). Cell replacement therapy *via* transplantation of mDA neuron precursors presents a promising strategy to reverse motor dysfunction in PD both at the cellular and circuit levels(3–5). Human pluripotent stem cells (hPSCs), comprising both human embryonic stem (ES) and induced pluripotent stem (iPS) cells, represent an attractive cell source for generating defined, scalable, and engraftable populations of mDA neurons(6–8). Grafting hPSC-derived mDA neurons has shown success in preclinical animal models of PD(9–12), and based on these results, cell replacement efforts using clinical grade mDA neuron precursors have moved from the preclinical phase to actual clinical translation in PD patients(13–17). Despite such rapid progress, one remaining challenge is the rather poor yield of mDA neurons, with typically around 10 % of total cells within the graft expressing tyrosine hydroxylase (TH)+, the rate-limiting enzyme for dopamine synthesis(13–16, 18, 19). In most studies, hPSC-derived mDA precursors yield a mixture of cells *in vivo* that includes non-mDA lineages such as subthalamic and hindbrain neurons, astroglia and oligodendroglia lineages and non-neural cells such as vascular leptomeningeal-like cells (VLMCs) within the graft(7). Thus, achieving an increased yield of authentic mDA neuron and a decreased proportion of potential off-target cell types remains an important goal.

Following early work demonstrating the derivation of mDA neurons *via* an hPSC-derived midbrain floor plate intermediate(20, 21), several groups have further optimized mDA neuron differentiation protocols by modifying timing and duration of patterning factors such as activation of canonical WNT, SHH, and FGF8 signaling. While all protocols transit through a midbrain floorplate intermediate, there is considerable variability in the yield of mDA neuron and the type and numbers of potential off-target populations across protocols(7, 22). The markers described for defining mDA neuron lineage such as FOXA2+ and LMX1A+ remain a critical component in the quality control of hPSC-derived mDA neuron products. However, they are not exclusive for mDA neuron identity as they can be expressed by precursors of the subthalamic lineage(12, 23). EN1 expression, in addition to FOXA2+ and LMX1A+, can help in better predicting successful graft outcome (12). During early midbrain development, EN1 *versus* DBX1 expression demarcate the midbrain *versus* the more anterior, diencephalic domain respectively(24). However, EN1 expression extends posteriorly into the anterior hindbrain. Thus, OTX2 expression, marking both midbrain and forebrain but not hindbrain anlage, can be used to further define midbrain floor plate identity and to mark the desired precursor population by quadruple expression of FOXA2+ LMX1A+ EN1+ and OTX2+. Previous work reported that late administration of FGF8b during mDA differentiation leads to improved induction and / or maintenance of EN1 among FOXA2+ and LMX1A+ precursors, and reduces anterior off-target cells, including glutamatergic neurons of the subthalamic nucleus(12). However, FGF8b treatment is also correlated with an increased proportion of VLMC-like cell types, expressing fibroblast markers such as COL1A1 and PDGFR *in vitro* and after *in vivo* engraftment (25, 26). Another study reported the manipulation of WNT, SHH, liver X receptor signaling to enhance mDA neuron yield while reducing off-target cell types. However, neither of those studies(27–29) performed an in-depth characterization of cell type identity, faithfulness of molecular marker expression and overall cell type composition *in vitro* and *in vivo*.

Postmitotic mDA neurons express Pitx3 and the deletion of Pitx3 results in the selective loss of mDA neuron in the substantia nigra (SNc)(30, 31). The derivation of SNc A9 mDA neuron subtype is particularly important for cell-based therapy in PD given the selective loss of A9 mDA neurons in PD, in contrast to A10 mDA neurons in ventral tegmental area that are relatively spared(2). Studies using scRNA-seq during mouse midbrain development identified ALDH1A1 as an important A9 mDA neuron marker(32) within the SOX6+ lineage. As such, ALDH1A1 is expressed during development and in adults in a subset of SNc mDA neurons that are highly vulnerable in PD(33). In addition, deletion of Aldh1a1 expressing cells in SNc in mice results in impaired motor skill learning (34). None of the current hPSC-differentiation protocols has demonstrated the molecular characterization of ALDH1A1+ A9 mDA neurons in hPSC-derived grafts, and the factors required to improve ALDH1A1 yield from hPSCs *in vitro* or upon transplantation *in vivo* remain unknown.

The increased use of single cell transcriptional analyses offers novel opportunities for defining cell type identity. Several recent studies characterized mDA neuron diversity during development *in vivo*(23, 32, 35, 36). However, there is a paucity of single cell data from hPSC derived mDA neurons after grafting. Early studies show that single cell RNA-sequencing of hPSC-derived mDA precursor grafts captures a very low yield of mDA neurons, representing less than 7% of the total cells sequenced, and comprising more than 90% off-target cells within the graft including VLMCs(26). Furthermore, the mDA neuron proportion present in the scRNA-seq study seems to differ significantly (up to 10-fold) from the percentage of mDA neurons reported in the corresponding histological data(26). This discrepancy may reflect the low-capturing efficacy of neurons by scRNA-seq due to technical challenges such as selective loss during enzymatic and mechanical dissociation(37). An improved understanding of cell type identity of hPSC-derived mDA cell products, including in-depth analysis of mDA neuron subtypes and off-target cell types, is essential to facilitate the prediction of graft success and reduce the risk of unwanted side-effects.

In the current study, we develop a novel protocol called “Boost+” that is based on our recently reported “Boost” differentiation conditions(9) which were the foundation of our clinical mDA product. Boost+ includes additional manipulations of FGF and WNT signaling at the stage of neurogenic differentiation, following midbrain floorplate induction. We apply FGF18 to highly maintain expression of EN1 under conditions that do not significantly induce VLMC-like lineages, and we concomitantly treat cells with the WNT inhibitor IWP2 to enhance the specification of the ALDH1A1+ lineages. The resulting Boost+ mDA cell product yields a higher portion of TH+ mDA neurons *in vitro*, and elevated expression of EN1 and PITX3. Furthermore, Boost+ patterned mDA neurons exhibit improved functionality based on electrophysiological properties and increased dopamine release. Single nucleus RNA-sequencing (snRNA-seq) demonstrates the presence of a highly enriched mDA neuron population within Boost+ grafts, compared to Boost grafts, allowing a detailed characterization of the molecular identity of mDA neuron subtypes and neuronal and non-neuronal contaminants. We observe a higher portion of mDA neuron population within the graft and a larger portion of ALDH1A1+ A9 mDA neurons, which also more closely resemble primary human fetal A9 mDA neurons (25, 26). Our *in vivo* functional studies indicate that both Boost and Boost+ grafts rescue drug and non-drug induced PD behaviors in the 6OHDA lesion rat model of PD, and that Boost+ graft exhibit recovery in additional non-drug-based behavioral assays such as ladder rung test. The Boost+ protocol improves the efficiency and specificity of mDA neuron differentiation from hPSC for application in both PD cell therapy and human disease modeling.

## RESULTS

### FGF18 and IWP2 at neurogenic conversion induce improved yield of authentic mDA neurons

We recently reported on a protocol to derive mDA neuron from hPSC suitable for clinical translation that involves a bi-phasic activation of WNT signaling: intermediate levels of WNT signaling to pattern neural identity towards midbrain fate, followed by the induction of high levels of WNT signaling mimicking the strong developmental expression of WNT1 at the anterior border of the midbrain hindbrain boundary. Under those “boost” conditions, we observed increased EN1 levels at day 11 compared to the same protocol without boost(9). We further demonstrated that EN1 is functionally important for mDA neuron specification as *EN1*-knockout lines show increased levels of forebrain and subthalamic markers compared to wild-type (wt)-hPSCs(9). However, despite increased levels at day 11, EN1 tends to decrease by day 16 of differentiation (**Fig. S1A**). In contrast, EN1 expression during normal midbrain development remains high within mDA neuron lineages, and EN1 expression is functionally important for mDA neuron survival(38–40). Loss of function studies in the mouse further indicate that FGF signaling is required to maintain EN1 expression that is necessary to induce mature DA neurons(41). Several groups have used FGF8b treatment during either early or late stages of mDA neuron induction and differentiation(12, 20). In the Boost protocol, exogenous FGF8b is not required for initial induction of EN1 at the floorplate intermediate stage, as the WNT-Boost triggers robust expression of endogenous FGF8b and FGF8-dependent genes such as *PAX2*, *PAX5*, *PAX8*(9). In contrast, late FGF8b treatment during neurogenic differentiation stage promotes maintenance of EN1 expression (**Fig. S1B, S1C**) but also induces markers of contaminating cell lineages such as SIX1 and SMA, a finding highly dependent on the onset and duration of FGF8b treatment (**Fig. S1B, S1C**). To overcome the trade-off between reduced EN1 maintenance and increased levels of SIX1 and SMA, we examined alternative FGF molecules among related FGF ligands, such as FGF8a, FGF17, and FGF18(42). The ligands FGF8a, FGF17 and FGF18 are selectively expressed in the midbrain after establishment of the midbrain/hindbrain organizer, while FGF8b expression becomes progressively restricted to the hindbrain(43). We found that FGF18 treatment can robustly maintain EN1 expression at day16 and 40 of differentiation while largely preventing the induction of SIX1 or SMA as compared to FGF8b treatment (**Fig. S1D, S1E**).

Next, we sought to further manipulate WNT signaling, a pathway crucial for mDA neuron specification(44). While WNT activation is essential for the robust induction of EN1 and suppression of subthalamic and forebrain fates, extended canonical WNT signaling increases the midbrain progenitor pool but results in incomplete mDA neurogenesis(45, 46). Accordingly, protracted periods of canonical WNT signaling may interfere with mDA neuron specification, and abrogation of WNT activation may be important for efficient mDA neuron generation including the derivation of A9 ALDH1A1+ mDA neuron. To address a role for late WNT inhibition, we conducted studies using the porcupine inhibitor IWP-2, which blocks canonical and non-canonical WNT signaling. Flow analysis, immunofluorescent staining, and RT-qPCR demonstrate that adding both IWP2 and FGF18 during neurogenic conversion under Boost conditions (called Boost+) leads to a high percentage of quadruple expression of FOXA2+LMX1A+OTX2+EN1+ at day16 (**Fig. 1A-1C, S1F, S1G**). Moreover, co-treatment of FGF18 and IWP2 induces increased A9 mDA marker (ALDH1A1) while further reducing off-target markers compared to FGF18 treatment alone (**Fig. S1G**). The Boost+ protocol is similarly effective for mDA differentiation in other hPSC lines, such as MEL1 hESC and control iPSC (J1) (**Fig. S1H, S1I)**. At postmitotic neuron stages, mDA neurons derived under Boost+ conditions showed increased biochemical markers of identity and mDA neuron fate, such as EN1, PITX3, and DAT (**Fig. 1D-1F**). Quantitative RNA FISH confirmed higher expression of *PITX3* mRNA expression in individual mDA neurons derived from the Boost+ *versus* Boost protocol (**Fig. 1E**).

**Figure 1.**
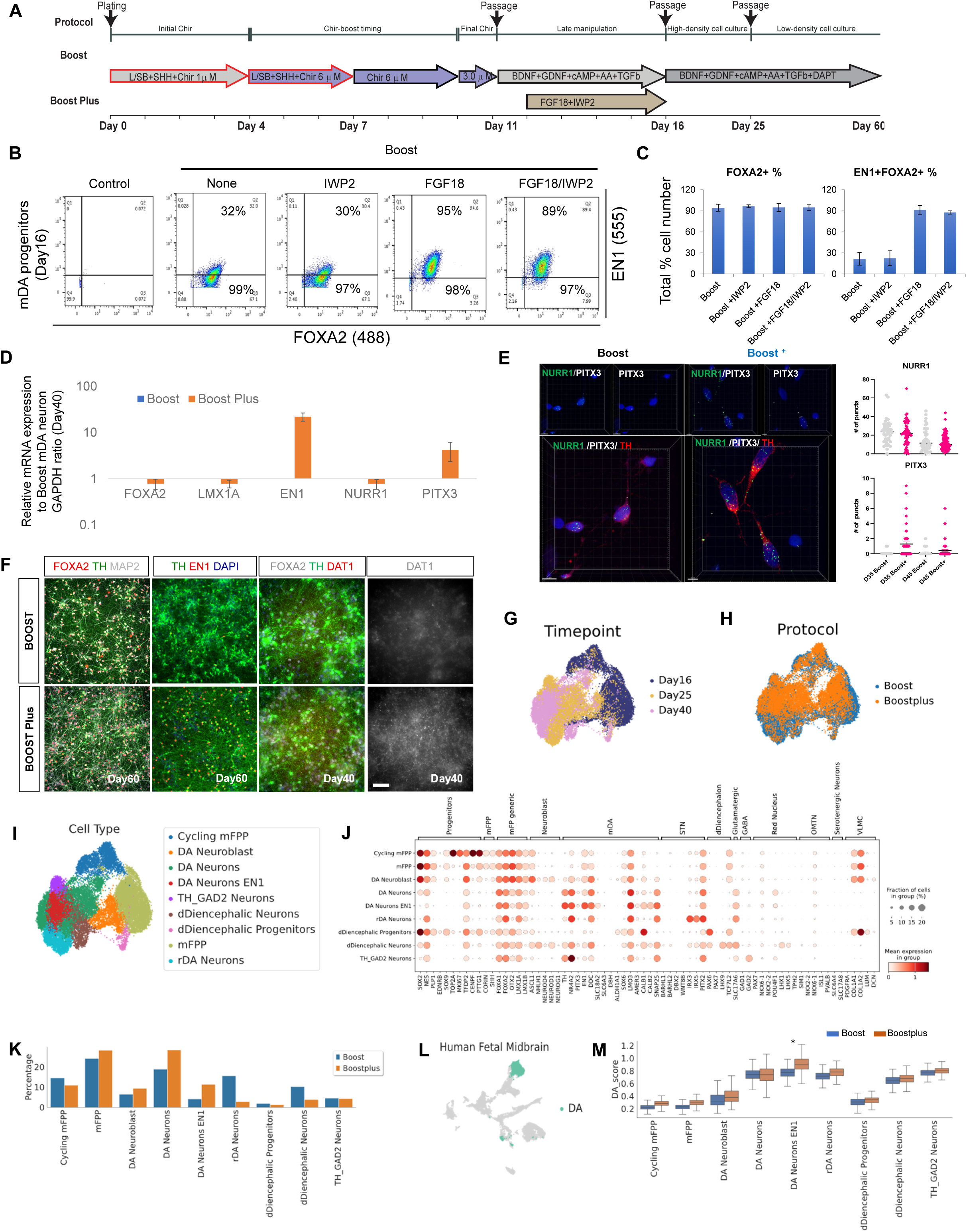
FGF18 and IWP2 at neurogenic conversion induce improved yield of authentic mDA neurons. **(A)** Schematic illustration of the Boost and Boost+ mDA differentiation method. **(B)** Representative dot plots from intracellular FACS flow analysis of day 16 differentiated progenitors patterned using various differentiation conditions for floor plate marker FOXA2 and midbrain marker EN1. On top of the Boost protocol, IWP2 alone, FGF18 alone, and IWP2+FGF18 were treated from day12 to day16. **(C)** Bar graph representing quantification for single positive (% FOXA2+) or double positive (% FOXA2+ EN1+) for each protocol (Boost, Boost + IWP2, Boost + FGF18, Boost + IWP2 + FGF18) (n=3) from **(B)**. **(D)** qRT-PCR analysis of mDA neurons at day40 for mDA neuron markers derived from the Boost and Boost+ methods. **(E)** Representative confocal image of RNA FISH of mDA neurons at day45 co-labeled with mature mDA markers (**left**). Human RNA probes against *NURR1* and *PITX3* were used with simultaneous protein staining of TH. Two different protocols were compared (Boost *vs*. Boost+). The number of RNA FISH dots for each RNA gene probe *NURR1* and *PITX3* were quantified among TH positive mDA neurons at different time points of differentiation (day35 and 45. **right**). **(F)** Immuno-fluorescent staining for mDA neuron markers, FOXA2, TH, EN1, and DAT1 in mDA neuron at day40 and day60 *via* the Boost and Boost+ conditions. **(G-I)** UMAP plots of *in vitro* scRNA-seq data colored by timepoint **(G)**, protocol **(H)** and cell type **(I)**. **(J)** Dotplot of canonical marker genes of midbrain progenitors, DA neuron subtypes and off-targets. **(K)** Overall percentage of cell types in each protocol. **(L)** UMAP of fetal midbrain dopamine (DA) neurons(69) selected to determine highly specific markers. **(M)** Boxplots showing the distribution of enrichment scores for each cell type and protocol according to fetal dopamine neurons. Mann-Whitney rank test and Benjamini-Hochberg correction; ∗p < 0.001.

To further assess cell type composition and molecular characteristics of the resulting cells, we conducted single cell RNA-sequencing (scRNA-seq) on days 16, 25, and 40 of mDA neuron differentiation comparing Boost *versus* Boost+ conditions **(Fig. 1G, 1H**). We classified cell clusters using canonical markers of neuronal progenitors and neurons of the ventral midbrain and surrounding regions (**Fig. 1I, 1J**, **S2A**). We observed that, at day16 of differentiation, most clusters represent ventral floorplate progenitors (FPP; **Fig. S2B**). At day 25, there was a mixed population of neuroblasts and neurons (**Fig. S2C**) while at day 40 of differentiation, we observed more mature neuronal lineages (**Fig. S2D**). Within the mature neuron clusters, mDA neurons were defined by the expression of *TH*, *NR4A2,* and low levels of *PITX3 expression,* and a subset of these neurons also expressed *EN1* (**Fig. 1J**). The population of mDA neurons expressing *TH*+ *EN1*+ increased by about 3-fold in the Boost+ condition (from 15% to 47%) by day 40 (**Fig S2D**) and showed expression of the dopamine transporter, *SLC18A2* and *SLC6A3* (**Fig. 1J, 1K, S2A**). Conversely, a rostral DA population (rDA neuron) expressing *TH*, *NR4A2*, *IRX2*, *IRX5* and *PITX2* (23) was also identified, representing a population enriched in the Boost protocol and relatively depleted in Boost+ (**Fig. 1J, 1K, S2A, S2C, S2D)**. A small proportion of off-target cells were also identified including population expressing dorsal diencephalic (dDiencephalic) markers compatible with pre-thalamic (*PAX6*, *LHX2*, *LHX9*, *LEF1*, *TCF7L2*, *SLC17A6*)(47–51) and pretectal neuronal fates (*MEIS2*, *LHX1*, *BARHL2*, *TBR1*)(48, 50, 52). Furthermore, we also observed a small population of *TH* and *GAD2*expressing cells that could represent an immature version of the *TH*^+^*GAD2*^+^*EBF2*^+^*CALCRL*^+^ population (**Fig. 1J, S2A, D**) recently discovered(53) within the adult primary midbrain. Finally, we could not identify any major representation of neurons of the subthalamic nucleus (STN), red nucleus, oculomotor neurons (OMNT). VLMCs were also not found as *PDGFRA* expression was not detected in combination with *COL1A1* and *COL1A2* (**Fig. 1J, S2A**). Strikingly, many specific lineage markers did not neatly segregate into a single cluster, with some cells co-expressing distinct lineage markers. For example, spurious expression of mDA neuron markers (*TH*, *NR4A2*, *EN1)* with a subthalamic marker *PITX2* or co-expression of *TH*, *NR4A2* with *POU4F1*. These hybrid states may be a result of the *in vitro* environment that triggers a cellular stress pathway as previously shown in organoids(54), which may interfere with complete lineage segregation. Nevertheless, the Boost+ protocol increases the yield of mature *TH*^+^ and *EN1*^+^ mDA neurons by three-fold and reduces the number of rostral DA neurons and diencephalic off-targets from 22% in the Boost to 8% in the Boost+ (**Fig. 1K**). Independent quantification of the number of cells co-expressing *EN1* and *TH* also confirmed a higher yield of mDA neurons in the Boost+ *versus* Boost protocol (**Fig. S2E**). Finally, to understand how mDA neurons authenticity differs between the Boost and Boost+ protocols, we scored the cells generated *in vitro* against a panel of 100 highly specific genes that characterize human fetal mDA neurons(55) (**Fig. 1L**). We observed that the highest score to primary fetal mDA cells was assigned to the *TH*+ *EN1*^+^ neurons of the Boost+ protocol, which show a significant increase in similarity compared to the Boost counterpart (**Fig. 1M**), indicating that this condition yields both a higher percentage but also more authentic mDA neurons compared to the Boost method..

### Improved dopaminergic function of Boost+ derived mDA neuron *in vitro*

We first measured the emergence of spontaneous *in vitro* network activity in mDA neurons derived from the Boost and the Boost+ conditions, using a high-density micro-electrode array (MEA) containing 4,096 electrodes and a 18kHz system. The Boost+ derived mDA neurons show higher levels of spontaneous activity than mDA neuron under the Boost conditions (**Fig. 2A**). In addition, high-performance liquid chromatography (HPLC) analysis of neurons at day 60 in the Boost+ protocol showed an 8-10 fold increase in dopamine levels upon KCI or Ca2+ stimulation, both in intracellular and dopamine release assays (**Fig. 2B**). Given the 2-3-fold higher percentage of TH+ mDA neurons in the Boost+ protocol, these data suggest higher levels of dopamine synthesis and release on a per cell basis when compared to Boost patterned mDA neurons. Next, basal membrane electrophysiological properties of mDA neurons (Boost+ *versus* Boost) were examined with patch-clamp recording on day 60. There were no major differences between the protocols, though Boost+ derived mDA neurons showed a trend towards lower input resistance, higher cell capacitance, and an increased spontaneous action potential (sAP) frequency (**Fig. 2C-2F**). Upon current injections, mDA neuron from Boost+ protocol showed an increased sAP frequency upon depolarization, maintained stimulated APs for longer time, and had a higher fraction of responding neurons than in the Boost protocol (**Fig. 2G-2I**). Taken together, there data suggest that mDA neurons under Boost+ conditions exhibit more mature functional properties using both electrophysiological and biochemical assays.

**Figure 2.**
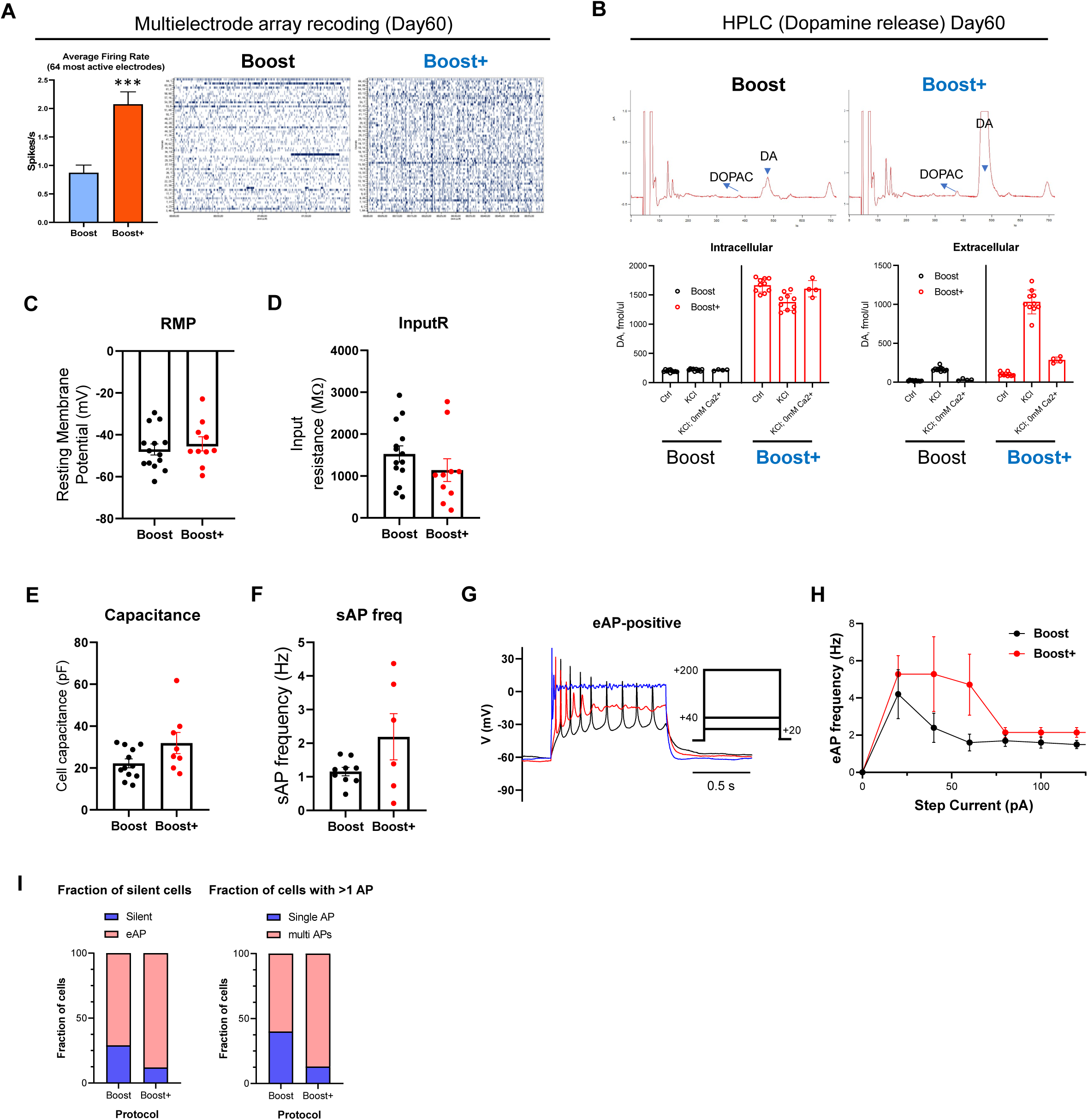
Improved dopaminergic function of Boost+ mDA neuron *in vitro.* **(A)** High-density multielectrode array recordings reveal increased firing rates at day50 mDA neuron derived from Boost and Boost+ protocols. **Left**, mean firing rates calculated from 60s of activity in the 1/64 most active electrodes of each probe (n = 256 electrodes from 4 MEA probes). **Right**, representative spike raster-gram displaying 1 m MEA recordings in Boost and Boost+ dopamine neurons. Data are represented as mean±SD. *** P<0.001. (**B)** Representative traces (**top**) and statistical analysis (**bottom**) of HPLC recording of DA release evoked by 80 mM KCl stimulation (5 min). Stimulation in the absence of extracellular Ca^2+^ is used as control. ***- p<0.001 from all other groups by 1-way ANOVA (n=10 culture dishes for control and KCl groups and n=4 for KCl in 0mM Ca^2+^). **(C-F)** Electrophysiological characteristics of cultured mDA neurons, including resting membrane potential (**C**, n=14 and 10 cells), input resistance (**D**, n=14 and 10), membrane capacitance (**E**, n=12 and 8) and spontaneous AP frequency (**F**, n=10 and 6). **(G)** Examples of mDA neuron responses to current injections. **(H)** Dependence of spike frequency evoked by current injection (eAP) on the current amplitude. ***- p<0.001 by two-way ANOVA. **(I)** Fraction of neurons responding to current injections was higher in Boost+ protocol (p<0.001 by χ^2^ test).

### *In vivo* cell type composition by single nucleus sequencing

To assess how the Boost+ protocol changes the cell type composition *in vivo*, we transplanted the same number of day16 mDA precursors patterned either *via* the Boost or Boost+ protocol into the striatum of adult NGS mice. At 1 month post implantation, we observed that the Boost+ grafts showed an increased yield of mDA neurons, expressing EN1 and ALDH1A1 by histological analysis while expressing comparable levels of FOXA2 between protocols (**Fig. 3A**). To define differences in cell type composition in more detail, we performed single nucleus sequencing (snRNA-seq) of the grafted Boost *versus* Boost+ lineages that stably express Td-Tomato at 1 month after transplantation (**Fig. 3B, S3A**). We used snRNA-seq as this technique should improve the efficiency of capturing neurons versus non-neural cells compared to scRNA-seq analysis(37). The snRNAseq data showed that grafted, unlike *in vitro* culture conditions, mDA neurons in the graft exhibit more conclusive mDA neuronal subtype identities including A9 mDA neurons expressing *ALDH1A1*, *SOX6* and *LMO3*, A10 DA neurons expressing *CALB1* and *CALB2* (**Fig. 3C, 3D**), and rostral DA neurons expressing low levels of *PITX2* together with *TH* and *NR4A2*. All mDA neurons expressed the canonical mDA markers including *TH*, *NR4A2*, *PITX3*, *EN1* and *DDC* in combination with the dopamine transporters *SLC18A2* and *SCL6A3* (**Fig. 3C, 3D**, **S3B**), implying a higher degree of maturity in grafted mDA neurons compared to the matched neurons *in vitro*. Beyond the large proportion of mDA neurons, we also defined a small percentage of off-target neuron types in the graft. In particular, we observed a subset of thalamic and pre-tectal neurons that were characterized by *LHX2*, *LHX9*, LEF1, *TCF7L2*(47–49) and *LHX1*, *LHX9*, *MEIS2*, *BARHL2* co-expression(50–52), respectively. An inhibitory sub-group of neurons was also defined, with a profile that resembled ventral lateral geniculate nucleus (dLGN) interneurons of the thalamus, with co-expression of *GAD1*, *GAD2*, *DLX1*, *DLX5*, *DLX6*, *OTX2* and *TLE4* (56, 57) (**Fig. 3C, 3D, S3B**). We observed that the transplanted cells for both Boost and Boost+ protocol yielded more defined neuronal subtypes with clearly distinct cluster and gene expression profiles as compared to mDA neuron cultures *in vitro*, consistent with the published literature that grafting organoid-derived cells *in vivo* can “rescue” limitations in cell subtype specification observed in organoid cultures *in vitro*(54). Partition-based graph abstraction (PAGA) analysis to understand the relationship between progenitors and neuron types demonstrated a link between mFPP and mDA neurons and an association between dorsal diencephalic progenitors and related off-target diencephalic neurons, suggesting these progenitors may be the ones that give rise to the diencephalic fates (**Fig. S3C**). There was no clear evidence of cells that recapitulate the identity of red nucleus neurons and OMNTs. Importantly, no VLMCs were detected that co-express *PDGFRA*, *COL1A1* and *COL1A2* (**Fig. 3D, S3B**). Similar to the *in vitro* results, the Boost+ protocol suppressed off-target diencephalic lineages to a higher degree (from 18% in the Boost to 5% in the Boost+) and enhanced the generation of mDA neurons from 46% in the Boost to 82% in the Boost+ (**Fig. 3E**). Importantly, the Boost+ protocol yielded more overall DA neurons including about twice the percentage of highly specific A9 mDA neurons (**Fig. 3E, 3F**), the population most critical for cell replacement therapy and recovery of motor dysfunction in PD. To independently validate the higher proportion of A9 mDA neurons in the Boost+, we sorted our snRNA-seq dataset based on *TH* and *NR4A2* and subclustered these cells to identify subgroups (**Fig. S3D, S3E**). Those data confirmed a higher proportion of *ALDH1A1* positive neurons in the Boost+ *versus* the Boost protocol (**Fig. S3F, S3G**). KEGG analysis revealed that the *ALDH1A1* compartment is enriched for dopamine signaling pathways while *CALB1* positive neurons show enrichment for morphine addiction, indicating a close relationship between specific gene signatures and the eventual function of these subtypes(58) (**Fig. S3H**). As both Boost and Boost+ protocols successfully differentiated into A9 mDA neurons upon transplantation, we next sought to identify any potential differences between these two A9 mDA populations (clusters 0 and 1; A9-like cells). Our analyses suggested that A9 DA neurons obtained *via* the Boost+ protocol exhibited significantly more elevated expression levels of TH, compared to those derived from the Boost protocol. Gene Ontology (GO) enrichment analysis further revealed that the Boost+ A9-like cells showed an upregulation of genes associated with glycolysis and ATP metabolism, implicating a potentially enhanced metabolic capacity in these cells (**Fig. S3I**). To test specific subtype authenticity, we compared our data to adult DA neurons within the ventral midbrain. First, we defined a highly specific signature of A9 (defined by *SOX6* positive neurons) and A10 (defined by *CALB1* positive neurons) mDA neurons by calculating the top differentially expressed genes in these two populations using a published dataset (59) (**Fig. 3G**). We then scored our cells against these two lists. We show that cells classified as A9 mDA neurons in our graft specifically enrich for *SOX6*+ A9 mDA neuron signatures (**Fig. 3H**), while grafted A10-like mDA neurons enrich for *CALB1*+ A10 adult mDA neuron signatures (**Fig. 3I**), confirming a match in identity of A9-like and A10-like mDA neuron in the graft to *in vivo* adult mDA neurons. Interestingly, when comparing scores between the Boost and Boost+ for authenticity, the Boost+ condition yields significantly more authentic A9 mDA neurons (**Fig. 3H**) while in the Boost protocol, the A10-like mDA neuron fraction showed higher authenticity compared to human adult A10 mDA neurons (**Fig. 3I**). Overall, we demonstrate that the Boost+ protocol increases the yield of A9 mDA neurons *in vivo*, with greater resemblance to adult primary A9 mDA neurons, and significantly reduces putative off-target diencephalic neurons.

**Figure 3.**
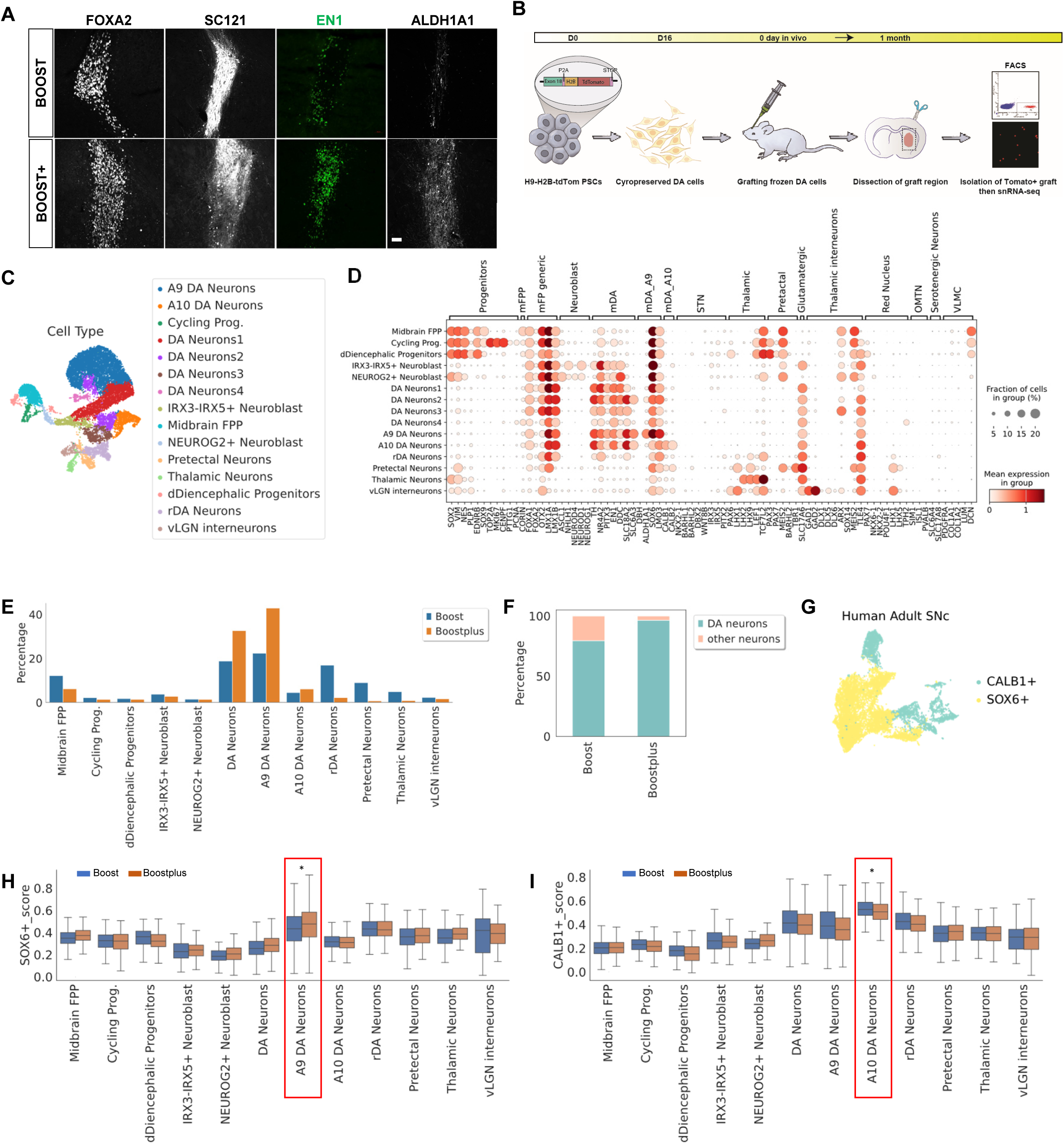
*In vivo* cell type composition by single nucleus RNA sequencing (snRNA-seq) **(A)** Representative microscopy images of 1-month-old (equivalent of day 45 differentiation *in vitro*) intrastriatal grafts from either Boost or Boost+ patterned progenitors (day16) on various makers, FOXA2, SC121, EN1, and ALDH1A1. Boost+ patterned neural precursors mostly maintained EN1 and ALDH1A1 expression *in vivo*. **(B)** Schematic illustration of the snRNA-seq from the grafts 1 month post implantation of the mDA neuron progenitor (day16) derived from the Boost and Boost+ method. **(C)** UMAP plot of 1 month grafted cells from Boost and Boost+ protocol. **(D)** Dotplot of canonical marker genes of midbrain progenitors, mDA neuron subtypes and off-targets **(F)** Percentage of mDA neurons in each protocol compared to other neuronal populations. **(G)** UMAP of adult midbrain dopamine neurons divided by SOX6 positive and CALB1 neurons (59) that were selected to determine highly specific markers for A9 and A10 DA neurons, respectively. **(H, I)** Boxplots showing the distribution of enrichment scores for each cell type and protocol according to A9 **(H)** and A10 **(I)** mDA neurons of adult signatures. Mann-Whitney rank test and Benjamini-Hochberg correction; ∗p < 0.001.

### *In vivo* functional characterization of Boost and Boost+ derived mDA precursor in 6-hydroxydopamine (OHDA)- induced Parkinsonian rats

Next, we grafted cryopreserved day 16 mDA neuron progenitors derived from either the Boost or Boost+ protocols into 6-hydroxydopamine (OHDA)-lesioned rats (immunocompromised NIH-*Foxn1^rnu^* strain) to evaluate graft survival and differentiation, and impact on behavioral deficits. 6-OHDA was unilaterally injected in the median forebrain bundle with subsequent amphetamine-induced rotation tests at 4 and 6 weeks (±1 week). Rats with more than six rotations per minute are defined as a complete lesion and were subsequently used in the grafting experiment. The selected 6-OHDA lesioned rats were randomized into three groups: vehicle, Boost cell, and Boost+ cell group. The Boost and Boost+ cells differentiations were performed on the same batch and were frozen on day 16 of differentiation. On the day of surgery, the Boost or Boost+ cells were freshly thawed and suspended at a concentration of 100,000 ± 10,000 cells/μl with a viability greater than 85%. Each animal received an injection of 400,000 cells into the striatum. Animals underwent a battery of behavioral tests at 0, 1.5, 3, 4.5, and 6 months (±1 week) post-grafting, including amphetamine-induced rotation, ladder rung walking test, and adhesive removal test. Both Boost and Boost+ cell grafts led to complete recovery at 4.5 months post-grafting in the rotation test (**Fig. 4A**). At 6 months after transplantation, the average of rotations per minute in Boost and Boost+ cell grafting group reached -2.68 and -3.75 respectively, compared with 10.65 in the vehicle group. In the adhesive removal test, both Boost and Boost+ cell groups showed significant recovery compared with the vehicle group at 6 months post grafting (**Fig. 4B**, left). Interestingly, only the Boost+ cell group showed significant recovery in the ladder rung walking test, compared with the vehicle group (**Fig. 4B**, right). Histological analysis of the brains showed that grafts from both Boost and Boost+ conditions survived and differentiated into TH^+^ mDA neurons (**Fig. 4C-4F**) and extending TH+ processes into the surrounding host tissue (**Fig. 4C-4F, S4A**). Stereological cell counts showed a significantly higher number of human cells (hNA^+^) in the Boost compared to the Boost+ grafts (**Fig. 4G**). The estimated graft volume in the Boost group (3.16 ±0.90 mm^3^) was also higher than that of Boost+ group (1.12±0.41 mm^3^), which is consistent with the human cell number. In contrast, the overall number of TH^+^ cells was similar in both groups (**Fig. 4G**). Accordingly, the proportion of TH+ dopamine neurons was significantly higher in the Boost+ graft reaching a mean of 30.8%, vs 9.4% in the Boost group (**Fig. 4H**), which is similar to the portion of TH ^+^ mDA neuron observed in the snRNA-seq at 1 month (**Fig. S3C**). Consistent with a higher percentage of TH+ cells, the density of TH expression was also higher in the Boost+ grafts than in the Boost grafts (**Fig. 4C-4F, S4A**). The vast majority of both Boost (average at 97.7%) and Boost+ (average at 96.7%) human cells express FOXA2 (**Fig. 4C, 4I**), but the percentage of EN1^+^ human cells was higher in the Boost+ grafts, (average at 47.6%) compared to the Boost grafts (average at 27.4%) (**Fig. 4D, 4I**). The percentage of EN1^+^ in hNA^+^ TH^+^ mDA neurons is also higher in Boost+ (average at 17.9%) than Boost cell grafts (average at 9.0%) (**Fig. 4D, 4J**). We also analyzed the grafts for GIRK2^+^ mDA neurons and found the percentage to be higher in Boost+ (92.9±1.7%) than Boost (59.7±6.8%) mDA neurons (**Fig. S4B**). In both Boost and Boost+ grafts, around 30% GIRK2 mDA neurons also co-express CALB1 (**Fig. S4B**). Expression of ALDH1A1, known to be expressed in mDA neurons in the human ventral SNpc (33), is also elevated in the Boost+ grafts (average of 26% of hNCAM^+^/TH^+^ human dopamine neurons) compared to the Boost (average at 3.3%) graft group (**Fig. 4E, 4J**). ALDH1A1^+^ dopamine neurons also extended axons into the surrounding host tissue (**Fig. 4E**). The vast majority (around 93%) of ALDH1A1^+^ dopamine neurons co-express GIRK2 in both Boost and Boost+ cell grafts (**Fig. S4C**). Dopamine transporter (DAT) expression was enriched in Boost+ cell grafts and more sparsely expressed in Boost cell grafts (**Fig. 4F**). To assess the proliferation rate of the grafted cells, we analyzed grafts for human specific Ki67 by IHC (**Fig. S4D**). The percentage of hKi67^+^ out of all human cells (hKu80^+^) was lower in the Boost+ (average at 0.7%) compared to the Boost (average at 2.8%) group (**Fig. 4K**), which may have contributed to the lower total human cell number in Boost+ than in Boost cell grafts (**Fig. 4G**). Taken together, these data suggest that that the Boost+ differentiation conditions led to a distinct graft composition with a greater proportion of mDA neurons, with a more A9-like profile, and a lower proportion of proliferative progenitors. We also performed IHC to detect non-neuronal cell types, including perivascular fibroblasts and choroid plexus-like epithelial cells, that were previously reported to be present in hPSC derived mDA cell grafts(11, 25), but were negative in our previously reported grafts(9, 14). Human specific hCOL1A1 labeled human perivascular fibroblasts, were negative in Boost grafts but present in Boost+ grafts (**Fig. S4E**). Transthyretin expressing cells, were absent in Boost grafts and sparse in Boost+ grafts (**Fig. S4F**).

**Figure 4.**
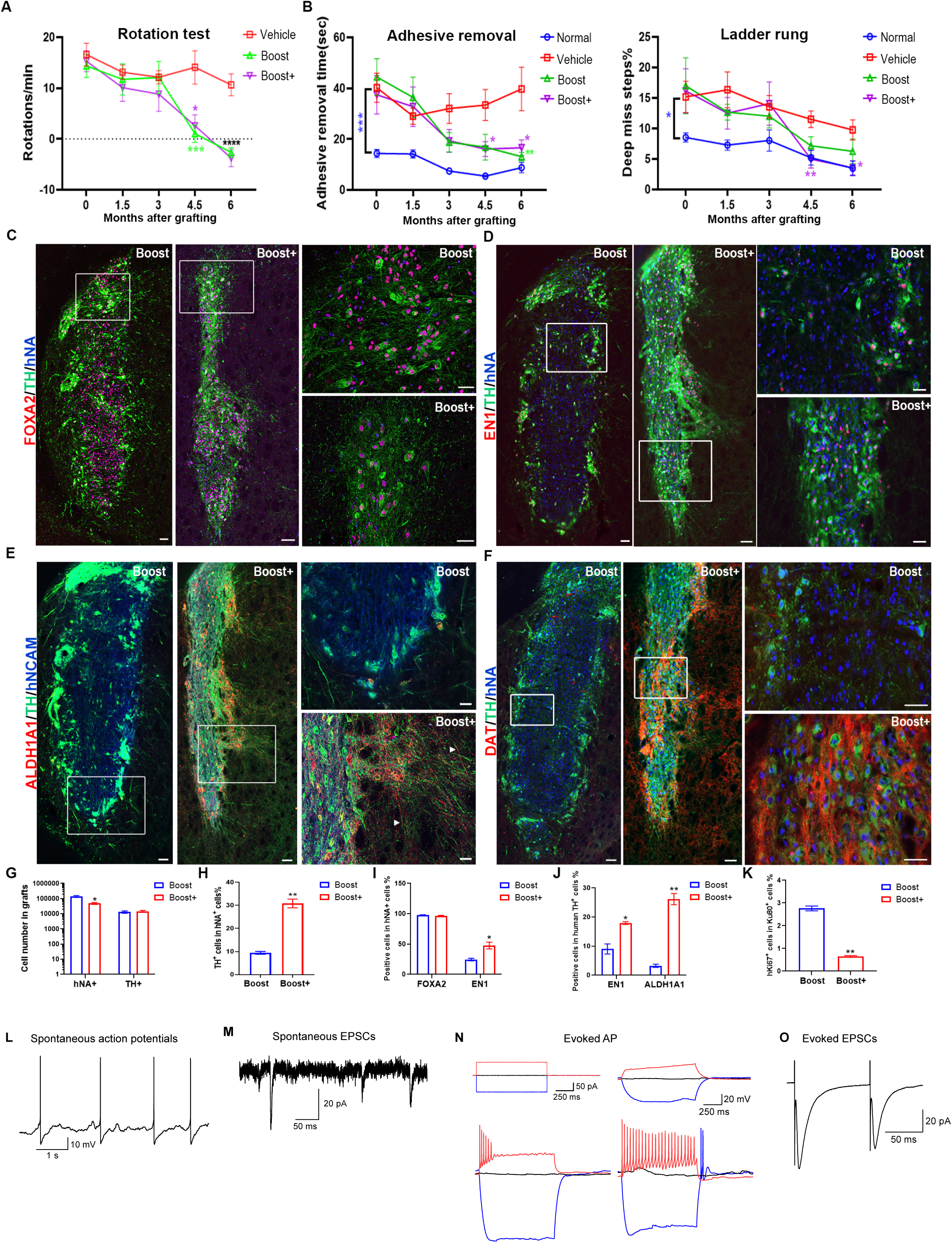
*In vivo* functional characterization of Boost and Boost+ derived mDA precursors in unilateral 6-hydroxydopamine (OHDA)-induced Parkinsonian rats. **(A)** Number of amphetamine-induced rotations per minute at several timepoints post-grafting in different animal groups. **(B)** The percentage of the adhesive removal time of the contralateral paw (**left**) and deep miss steps (**right**) in the ladder rung walking test at different time points after transplantation in different animal groups. **(C)** Representative images of immunohistochemistry (IHC) for FOXA2, tyrosine hydroxylase (TH) and human nuclei antigen (hNA) in Boost and Boost+ grafts. **(D)** Representative images of IHC for engrail 1 (EN1), TH and human nuclei antigen hNA in Boost and Boost+ grafts. **(E)** Representative images of IHC for ALDH1A1, TH and human specific marker hNCAM. The arrowheads point to TH ^+^ dopaminergic axons expressing ALDH1A1. **(F)** Representative images of IHC for dopamine transporter (DAT), TH and human nuclei antigen hNA in Boost and Boost+ grafts. Note that more DAT+ area is present in Boost+ than Boost grafts. **(G)** hNA ^+^ and TH ^+^ cell number in Boost and Boost+ cell grafts estimated by stereology calculation. **(H)** The percentage of TH+ out of hNA ^+^ cell number in Boost and Boost+ cell grafts. **(I)** The percentage of FOXA2 or EN1 expressing hNA ^+^ cell number out of hNA ^+^ cell number in Boost and Boost+ cell graft group. **(J)** Boost+ grafts have higher percentage of EN1+TH+ and ALDH1A1+TH+ cells out of the total human TH+ cells than Boost grafts. **(K)** The percentage of human specific hKi67+ cells out of human specific hKu80+ human cells in Boost and Boost+ cell grafts. Animal numbers are 5-7 per group in **A-B** and 3 per group in **G-K**. Data are represented as mean±SEM. *P<0.05, **P<0.01, ***P<0.001. In **A-B**, the color of * represents the significance between the group with the same color of symbol and the vehicle group. In **C-F**, scale bar represents 100 µm on the left panels while 50 µm on the right panels. The areas outlined in white lines are shown with higher magnification images on the right panels. **(L-O)** Whole-cell recordings from grafted mDA neurons (Boost+: 6 months post transplantation) including examples of a spontaneously active neuron **(L)**, spontaneous EPSCs **(M)**, action potentials evoked by current injections **(N)**, and EPSCs evoked by local electrical stimulation **(O)**. Recordings were done in the absence of synaptic blockers.

Finally, we performed whole-cell recordings from the graft derived mDA neurons from the Boost+ group at 6 months post implantation. We observed that the grafted neurons exhibit spontaneous activity, spontaneous EPSCs, action potentials evoked by current injections, and EPSCs evoked by local electrical stimulation (**Fig. 4L-4O**). In addition, whole-cell recordings from striatal spiny projection neurons (SPN) proximal to the mDA neurons graft indicate that both the Boost and Boost+ grafts mediate action potential frequencies back to levels comparable to recordings in non-lesioned mice (sham). In contrast, a lower AP frequency in SPN was observed when recorded in lesioned control mice (no grafts). However, only Boost+ graft triggered a significant reduction in resting membrane potential (RMP) and EPSC amplitude in SPNs upon current injection conditions (**Fig. S4G-S4N**).

## DISCUSSION

hPSC-derived mDA neurons hold great promise for cell replacement therapy in PD(3–5). Indeed, early-stage clinical trials of hPSC-based cell therapy in PD have been initiated in several countries(13–16, 60). To maximize the safety and efficacy of hPSC-based cell therapy approaches, differentiation protocols need to yield high percentages of authentic mDA neurons while minimizing off-target cell types. To reach this goal, unbiased methods are required for improved characterization of cell type composition in hPSC-derived mDA neuron lineages *in vitro* and after transplantation *in vivo*.

In this study, we present a novel protocol that we call Boost+, which includes the application of FGF18 and IWP2 at the stage of neurogenic conversion, built on top of the previously published Boost protocol. Boost+ improves the efficiency of midbrain patterning and mDA neurogenesis, with higher sustained expression of EN1 and the enhanced induction of authentic A9 and mature mDA neuron markers, such as ALDH1A1, PITX3, and DAT (**Fig. 1A-1F and S1F-S1I**). Systematic characterization by scRNA-seq *in vitro* confirms improved mDA neuron yield and reduced off-target cell types (**Fig. 1G-1K and S2**). Furthermore, Boost+ derived mDA neurons more closely match the transcriptional profile of *in vivo* human mDA neuron (**Fig. 1L, 1M**). *In vitro* functional assays, including MEA, HPLC, and electrophysiology, demonstrate that hPSC-derived mDA neurons derived *via* Boost+ show increased functionality at comparable differentiation stages (**Fig. 2**).

Additionally, we present snRNA-seq data that show highly mDA neuron enriched grafts *in vivo* derived from an hPSC-based cell product, a finding not seen in previous *in vivo* single cell studies of mDA neuron grafts where single cell results were dominated by non mDA neuron populations(25, 26, 35). Our snRNA-seq data enabled in-depth analysis of mDA neuron subtypes within the graft. Those data confirm that Boost+ condition yields a higher percentage of A9 mDA neurons expressing ALDH1A1, EN1, DAT, and TH, and that those neurons are more authentic to human A9 mDA neurons *in vivo* (**Fig. 3 and Fig. S3**). On the other hand, the snRNA-seq data also define novel types of neuronal contaminants including pretectal and thalamic neurons and show that those off-target-neurons are reduced in Boost+ *versus* Boost (**Fig. 3E, 3F and Fig. S3B, S3C**). *In vivo* functional studies in a preclinical PD rat model indicate that both Boost and Boost+ can rescue drug-induced rotation behavior and non-drug-based assays such as adhesive removal test. However, only Boost+ grafts induced significant recovery in the ladder rung walking assay (**Fig. 4A-4B**).

The low proportion of mDA neurons in the final graft composition (typically less than 10%) has been a challenge for the cell replacement therapy in PD(13–16, 18, 19). Purification using mDA cell product related surface markers or reporter lines have been employed to enrich mDA portion in the final graft(61–63). However, the use of reporter lines may not be suitable for clinical use, and using surfacer markers can also be challenging for the routine application in cell-based therapy by requiring clinical grade antibodies and sorting strategies. Furthermore, there is no consensus on the most appropriate surface markers for mDA neuron sorting. Our stereological quantification of the grafted cells shows that the Boost+ graft exhibits a higher portion of TH+ neurons (∼30%) out of total graft cells (**Fig. 4H**). This is a high TH density compared to other hPSC-derived mDA cell grafts without purification prior to the implantation steps. Interestingly, the proportion of TH+ mDA neurons in the final graft is similar to the proportion of TH+ cells from the snRNA-seq of Boost+ mDA cell graft at 1 month (**Fig. S3D**). Those results suggest a high consistency between snRNA-seq and histology results as well as high reliability of our snRNA-seq analysis; typically, snRNA-seq analyses of grafted cells show a drop-off of the neuronal population (25, 26).

Both Boost+ and Boost protocols yield a higher portion of ALDH1A1+ neurons in the 1-month grafts compared to the matched day 40 *in vitro* conditions. These results indicate that the *in vitro* culture conditions are less favorable for ALDH1A1 induction and maintenance compared to the *in vivo* striatal brain environment in the host brain. Indeed, the scRNA-seq analysis of *in vitro* maintained cells indicate that expression of the dopamine transporters *SLC18A2* and *SCL6A3* is lower than at matched *in vivo* stages in the graft, reflective of an immature mDA profile. A9 mDA neuron, particularly the ALDH1A1+ A9 mDA neuron subtype, is well-known to be highly vulnerable in PD(2). Therefore, it is advantageous to develop conditions that improve our ability to manipulate ALDH1A1+ A9 mDA neuron specification and maintenance in culture. Such conditions should enable improved disease modeling strategies and more precise cell-based therapy approaches, with highly enriched A9 mDA neuron grafts. Our data demonstrate a greater midbrain authenticity and maturity of *in vivo* grafted *versus in vitro* cultured mDA neurons at comparable stages of differentiation which may enable the identification of factors that drive more complete mDA neuron differentiation by comparing age matched *in vitro* scRNAseq and *in vivo* snRNAseq data (**Fig. S5**). The KEGG analysis indicate that Rap1 and GnRH secretion may show increased signaling and pathway activation in mDA neurons in the graft compared to the culture conditions, both in the Boost and Boost+ conditions (**Fig. S5**). Further studies will be required to determine whether those pathways and signals are involved in the maintenance and / or maturation of ALDH1A1+ A9 mDA neurons in culture.

The behavioral data support similar ability of the Boost and Boost+ mDA neurons to reverse drug-induced rotations and recover the task of adhesive strip removal. Interestingly, the ladder rung walking test is improved only in the Boost+ mDA group. This test is recognized for its greater sensitivity to subtle impairment in locomotor function (64) and tests motor function independently of the rotational score (65); it would therefore suggest a more complete recovery of locomotion by the Boost+ mDA cells. It will be interesting in future studies to determine whether Boost+ mDA neuron grafts can achieve recovery with lower numbers of grafted cells and whether the minimal threshold of surviving mDA neurons for functional recovery is different between Boost *vs* Boost+ mDA neurons.

In conclusion, our improved mDA differentiation method generates a higher yield of authentic functional mDA neurons and a lower number of cell contaminants in both cultured cells and in the resulting grafts by both histological and by scRNA- and snRNA-seq data. Therefore, the Boost+ condition may present a next generation of a scalable, off-the-shelf mDA cell product for clinical use in human. Such a product may offer improved therapeutic potential based on our *in vivo* grafting data, with highly enriched TH+ and ALDH1A1+ A9 mDA neuron proportions and with increased DAT expression. However, one of the limitations of the current study is the slightly worse performance of the Boost group as compared to the Boost group data observed in our preclinical batch(9), such as a slightly lower percentage of TH+ cells and a slight increase in Ki67+ cells at 6 months post grafting. Future studies will be required to confirm the relative magnitude of the approximately 3-fold increase in the percentage of TH+ cells under Boost+ *versus* Boost conditions observed in the current study. We have recently reported that grafting post-mitotic mDA neurons, purified *via* surface markers, such as low-49e and high-184 is feasible. Furthermore, we showed that adding the FDA approved TNF-alpha inhibitor, adalimumab greatly improves survival in such grafts(66). While those studies were performed using Boost mDA precursors, our current study suggests that cell sorting and adalimumab treatment may be particularly suitable in the Boost+ condition. Combining the Boost+ protocol with cell purification and adalimumab treatment could represent a strategy to achieve fully postmitotic and highly mDA neuron- and A9 subtype-enriched grafts for preclinical studies and potentially for eventual human translation in PD patients.

## MATERIALS and METHODS

### hPSC culture

Human pluripotent stem cells [hPSCs; WA09 (H9; 46XX) and MEL1 (46XY)], and J1 human induced PSC (MRC5), which was previously published in(67), were cultured onto Vitronectin (VTN-N, Thermo Fisher #A14700) coated dishes with Essential 8 media (Life Technologies #A1517001). Passage 35-55 hPSCs were used for the experiments. hPSCs were sub-cultured every 4-5 days by EDTA. All cell lines are cultured at 37°C with 5% CO2 and routinely tested for mycoplasma. All stem cell work was conducted according to protocols approved by the Tri-Institutional Stem Cell Initiative Embryonic Stem Cell Research Oversight Committee (Tri-SCI ESCRO).

### Transfection of hPSC

hESCs were dissociated to a single cell suspension with Accutase (Innovative Cell Technologies # AT104) and plated at a density of 250,000cells/well in a vitronectin coated 6-well plate in Essential 8 supplemented with 10 μM Y-27632 ROCK inhibitor (Bio-Techne #1254/50). The following day, media was replaced with 2ml mTeSR (STEMCELL Technologies #85850) supplemented with CloneR (STEMCELL Technologies #05888). Lipofectamine-DNA complexes were assembled in Opti-MEM (Thermo #31985062) according to the manufacturers protocol, using 8ul of Lipofectamine Stem (Thermo #STEM00001) and 5ug plasmid DNA per well. 24 hours post transfection, media was replaced with Essential 8 (Thermo #A1517001) supplemented with CloneR. 48 hours following transfection, hESCs were dissociated to single cells with Accutase and GFP+ cells that received the pX458 vector were isolated via FACS with a BD Aria6. Sorted cells were replated in vitronectin in Essential 8 with CloneR. CloneR was removed after 4 days according to the manufacturer’s protocol and clones were picked and assessed for either deletion or transgene incorporation.

### Generation of reporter lines

*GPI* targeting constructs were generated from a genomic DNA PCR with Q5 polymerase (New England Biolabs #M0494) amplifying ∼500bp of homology per side and assembled with NEBuilder (New England Biolabs #E2621S). An sgRNA targeting *GPI* (GPI sgRNA: CTTCATCAAGCAGCAGCGCG) was co-transfected with its respective targeting constructs for line generation. hESC lines were transfected as described above, for reporter lines, a ratio of 1:5 of sgRNA vector to targeting vector was used. Clones were screened *via* genomic PCR for the expected insertion.

### Directed differentiation into midbrain dopamine neurons from hPSC

hPSCs were dissociated into single cells using Accutase, and plated at 400K cells/cm^2^ onto Geltrex (Life Technologies, #A1413201) coated dishes with Neurobasal (Life Technologies)/N2(Stem Cell Technologies)/B27(Life Technologies) media containing 2mM L-glutamine, 500ng/ml SHH C25II (R&D systems #464-SH), 250nM LDN (Stemgent # 04-0074-02), 10μM SB431542 (R&D systems #1614), 1μM CHIR99021 (R&D systems #4432), and 10uM Rock inhibitor (Y-27632, R&D systems #1254), which represents day 0 of differentiation, and cultured until day 3 without Rock inhibitor from day 1. On day 4, cells were exposure to different concentration of CHIR 6 μM until day 10. On day 7, LDN, SB, and SHH were withdrawn. On day 10, media was changed to Neurobasal/B27/L-Glu supplemented with BDNF (brain-derived neurotrophic factor, 20ng/ml; R&D #248-BD), ascorbic acid (0.2 mM, Sigma #4034), GDNF (glial cell line-derived neurotrophic factor, 20 ng/ml; Peprotech # 450-10), TGFβ3 (transforming growth factor type β3, 1 ng/ml; R&D #243-B3), dibutyryl cAMP (0.2 mM; Sigma #4043), and CHIR 3 μM. On day 11, cells were dissociated using Accutase and replated under high cell density (800K cells/cm^2^) on polyornithine (PO; 15 μg/ml)/ laminin (1 μg/ml)/ fibronectin (2 μg/ml) coated dishes in mDA differentiation media [(NB/B27/L-Glu, BDNF, ascorbic acid, GDNF, dbcAMP, and TGFβ3 until day 16. For the improved protocol, Boost+, IWP2 (1uM, Tocris Bioscience #3533) and FGF18 (100ug/ml, Peprotech #100-28) were added from day12 to day16. On day 16, cells were dissociated and plated as same procedure of day 11 and cultured until day 25 using mDA differentiation media with adding DAPT (10 μM, R&D #2634)]. On day 25, cells were dissociated using Accutase and replated under low cell density (200K∼300K cells/cm^2^) in mDA differentiation media until the desired experiments. For the cryopreservation of mDA neuron precursor, day16 mDA differentiated cells were treated with Accutase, washing, detached, making single cells, and pelleting. Cell pellets were resuspended at a cell density of 8 million cells/mL of STEM-CELLBANKER. Controlled rate freezer (ThermoFisher) was used to cryopreserve cell product.

### Immunohistochemistry

Cells were fixed in 4% paraformaldehyde (PFA) (Affymetrix #MFCD00133991) in DPBS for 15 min at room temperature. Cells were subsequently washed with DPBS. Then cells were permeabilized with 0.5% Triton X-100 for 20min and blocked with 2% BSA in DPBS for 30min. The samples were subsequently incubated with primary antibody overnight at 4°C. The next day, after washing with DPBS, the samples were incubated with secondary antibody conjugated with Alexa Fluor 488-555-, or 647- (Thermo Fisher) diluted at 1:400 in 2% BSA (DPBS) for 1 hour at room temperature in shaking incubator. Next, the samples were washed with DPBS and count-stained with 4′, 6-diamidino-2-phenylindole (DAPI) (Sigma, #D9542). Images were visualized using an Olympus and Zeiss inverted fluorescence microscope. Mouse and chicken anti-MAP2 (1:1500, Sigma and 1:2000, Abcam), rabbit and mouse anti-TH (1:500, PelFreez and 1:1000, Immunostar), goat anti-FOXA2 (1:200, R&D), Rabbit anti-LMX1A (1:1500, Abcam), Goat anti-OTX2 (1:1000, Neuromics), rabbit and mouse anti-PAX6 (1:500, Covance and 1:200, BD-Biosciences), mouse and rabbit anti-EN1 (1:50, DSHB and 1:200 Invitrogen), goat anti-ALDH1A1 (1:250, Santa Cruz; R&D #AF5869), rabbit anti-GIRK2 (1:400, Almonte), rabbit anti-CALB1 (1:2000 Swant), and mouse anti-NURR1 (1:1500, Perseus Proteomics) were used for immuno-fluorescent staining. Donkey anti-mouse, goat, rabbit or chicken secondary antibodies conjugated with Alexa Fluor-488, Alexa Fluor-555 or Alexa Fluor-647 fluorophore (1:400, Life technologies) were used. Nuclei were counterstained by DAPI.

### RNA extraction and Real-time qRT-PCR

Total RNAs from samples were isolated with TRIzol (QIAGEN) using the Direct-zol RNA MiniPrep kit (Zymo Research, #R2052). 1ug of RNA was used to generate cDNA using the iScript Reverse Transcription Supermix (BioRad, #170-8841). Real-time qRT-PCR was performed using the SSoFAST EvaGreen Mix (BioRad) in a BioRad CFX96 Thermal Cycler. All reactions were performed according to the manufactured protocol. Primer sequences are listed below, and some primers were obtained from QIAGEN (Quantitect Primer assays). Results were normalized to GAPDH. Primer sequences are listed in Table S1.

### Multi-electrode array recording

A 100-μl droplet of medium containing 300,000 hPSC-derived midbrain dopamine neurons was seeded onto poly-l-lysine-coated complementary metal oxide semiconductor multi-electrode array probes (CMOS-MEA). 1.5 ml of medium were added after 1 h of incubation and replaced every 3 days. MEA recordings were performed 24 h after each medium change. 1 minute of spontaneous activity was sampled from 4096 electrodes using the BioCAM system and analyzed using BrainWave 4 software (3Brain AG, Pfäffikon, Switzerland). Spikes detection was performed using a Precise Timing Spike Detection (PTSD) algorithm with a detection threshold of 9 standard deviations.

### Single nuclei preparation

Nuclei isolation protocol was adopted from the previous study (68). The tissue region of interest, the striatum, was grossly dissected out using a pre-chilled scalpel on ice and minced into small chunks, and transferred using low-attachment 1,000p pipet by resuspending them in 1 mL of ice-cold homogenization buffer (HM). Tissue resuspension in a glass dounce homogenizer was gently homogenized with 10 strokes of the loose (A) pestle, followed by 20 strokes of the tight (B) pestle. The homogenized suspension was transferred to a pre-chilled low DNA-bind Eppendorf tube and centrifuged for 1,000g for 8 minutes at 4°C countertop microcentrifugation. The pellets were then gently resuspended in 250 µL HM buffer, which was then later mixed with 250 µL of 50% iodixanol mixture. During optiprep gradient system where 500 µL nuclei mixture was overlayed on top of 500 µL of 29% iodixanol solvent in a pre-chilled Eppendorf tube for high-speed centrifugation at 13,500g for 20 minutes at 4°C countertop microcentrifugation. The pellet is resuspended in nuclei storage buffer (NSB), counter stained for DAPI, and processed for sorting to enrich human nuclei.

The composition of each buffer was the following: HM is made from NIM2, 0.1% Triton-X 100, 40U/µL Rnasin, 20U/µL Superasin, and 5 µg /ml actinomycin D. Nuclei isolation medium 2 (NIM2) was made fresh for immediate use and discarded which is made up of NIM1, 0.5 mM DTT (dithiothreitol), and protease inhibitors (Roche mini complete, EDTA-free) 1 tablet for 10 mL solution. Nuclei isolation medium 1 (NIM1) was composed of 320 mM sucrose, 0.1 mM EDTA, 10 mM Tris buffer (pH 8.0), 5 mM MgCl_2_, 150 mM KCl, nuclease-free water (up to 100 mL). Iodixanol medium diluent (IDM) was made up of 250 mM sucrose, 150 mM KCl, 30 mM MgCl_2_, 10 mM Tris buffer (pH 8.0), and nuclease-free water (up to 100 mL). To prepare 29% and 50% iodixanol gradient, optiprep 60% iodixanol (D1556-250ml from Sigma) was diluted with IDM as solvent. Nuclei storage buffer (NSB) was 166.5 mM sucrose, 4 mM MgCl_2_, 10 mM Tris buffer (pH 8.0), 1 \% BSA, nuclease-free water (up to 100 mL), 5ug/ml actinomycin D, 40U/µL Rnasin, and 20U/µL Superasin.

### Single-molecule RNA fluorescent in situ hybridization (smFISH)

ViewRNA Plus ViewRNA™ Cell Plus Assay Kit (Invitrogen) kit was used in RNAse free conditions throughout the experiments. Adherent cells plated in a confocal-friendly plastic bottom µ-plate 24 Well Black (Ibidi) plates were fixed and permeabilized for 15 min at room temperature (RT) with Fixation/Permeabilization solution and blocked for 20 min followed by incubation with primary antibody TH; tyrosine hydroxylase and secondary antibody Alexa Fluor 647 (Invitrogen) for 1h at RT to locate RNA puncta signals within a mature DA neuron. Following protein detection, a fluorescent in situ hybridization (FISH) and branched DNA amplification technology is used to amplify the signal detection of an RNA transcript. ViewRNA Cell Plus assay (Invitrogen) protocol can address more detailed experimental steps.

Briefly, in the first step, a gene-specific oligonucleotide target probe binds to the target RNA sequence. Type 1 (Alexa Fluor 546) and type 4 (Alexa Fluor 488) probe sets were used to detect various genes. Pre-amplification and amplification steps were then achieved through a series of sequential hybridization steps at 40°C under gentle agitation for 1 h each. After two sequential amplifying steps, a fluorescent dye was introduced to hybridize to their corresponding amplifier molecules at 40°C under gentle agitation for 1 h. Washes were performed as indicated in the kit’s procedure. Samples were incubated with 5 µM DAPI to visualize cell nuclei and a coverslip was gently placed inside each well using ProLong™ Glass Antifade Mountant. RNA signals in dots were visualized by acquiring a series of z-stack images at 0.4µm step and covering the entire cell’s volume using SP8 Leica point scanning confocal microscopy with 63X oil lenses with optical zoom of 3. All the images in z-stacks were projected and obtained using Imaris software. Projected images were analyzed for quantification to localize the number of RNA molecules within each TH-positive dopamine neuron. Eight different fields of view (2-5 neurons/field) for each condition (boost vs. boost plus) from four independent differentiation batches (16 fields of view/condition) were obtained for downstream analysis. Gene targeting probes included human specific PITX3 and NR4A2. Human specific ViewRNA Cell Plus gene probes were designed by ThermoFisher https://www.thermofisher.com/order/genome-database/embeddedSearchForm?searchFormId=quantigene-viewrna-cell-only). PITX3-Alexa 647 Type 6 (VA6-3168220-VCP) and NR4A2-Alexa 488 (VA4-3082508-VCP) were used in the experiment.

### Intracellular protein staining and flow cytometry analysis

Cells were dissociated into a single cell using Accutase for 30 minutes at 37℃, washed with DMEM base medium (ThermoFisher), and filtered through the 30 µm blue cap strainer. Cells were then fixed with BD cytofix/cytoperm solution and incubated for 30 minutes on ice or at 4℃, followed by three washes with BD perm/wash buffer solution (cat. 554723). Cells were pelleted down at 500 g for 5 minutes at pre-cooled countertop microcentrifuge, resuspended, and incubated in an antibody cocktail diluted in BD perm/wash buffer at 4℃ or on ice for 30 minutes. For FOXA2 (R&D), OTX2 (R&D), and EN1 (Invitrogen), a primary antibody for intracellular stain dilution was employed 1 to 25 ratio. Upon the primary antibody conjugation, cells were washed three times with BD perm/wash buffer solution, followed by an incubation with a secondary antibody cocktail reaction (1 to 5,000 dilution in perm/wash buffer) at 4℃ for 30 minutes. Two additional washes with BD perm/wash buffer solution were performed with the final wash in 1x PBS. Samples were stored at 4℃ protected from light until FACS analysis. For flow analysis, FACSAria III flow cytometer (BD Bioscience) was used to record the expression pattern of single cells and FlowJo software was used to analyze between conditions. Proper isotype and secondary alone controls were used for gating.

### Tissue processing for immunofluorescence (IF) labeling

Histology on tissues from mice was performed on frozen sections from xenografts. Mice were anesthetized with pentobarbital and transcranial perfused using heparinized (10U/mL) PBS (pH 7.4), followed by 4% paraformaldehyde in PBS. The liquid was administered using a peristaltic pump to control the rate of the solution delivery to the system. Tissues were post-fixed in ice-cold 4% paraformaldehyde for 18 hours sharp and transferred to 30% sucrose until the tissue sink (typically 3-6 days post-), followed by snap freezing in O.C.T (Fisher Scientific, Pittsburgh, PA) or Neg-50 (Thermo Scientific). Brain tissues were all sectioned in 30 µm thick coronal sections using a cryostat and mounted onto a Superfrost plus microscope slides (Fisher Scientific). All the slides were air-dried for 18 hours at RT and stored at -80℃ for long-term use.

To process for immunolabeling, tissues were washed twice with 1x PBS, followed by permeabilization in 0.5% Triton X-100 in PBS for 10 minutes. Living cells in culture were directly fixed in 4% paraformaldehyde for 18 min, followed by 10 min permeabilization in 0.5% Triton X-100 in PBS. For labeling, cells or tissue sections were immunostained with primary antibodies of interest in 2% BSA in 0.25% Triton X-100 in PBS at 4°C overnight. The next day appropriate Alexa Fluor secondary antibodies were conjugated at room temperature for 30 minutes at a dilution of 1:500. Nuclei were counterstained by DAPI.

### Single cell analysis

The samples underwent 10X chromium Single Cell 3’ v3 processing. The reads were aligned to the human reference GRCh38 genome (and mouse mm10 in grafted cells) using Cell Ranger 5.0.0. The resulting filtered count matrix was further filtered for human cells in the grafted datasets. Downstream analyses were performed using the Scanpy v1.9.3^14^. All genes expressed in less than 5 cells were removed. In vitro data and graft data were processed separately, and quality control metrics were based on specific values in each dataset. Cells with less than 500 genes and more than 2000 genes were removed in the in vitro data, while for the graft data cells were kept within the range of 1000-5000 genes to control for doublets and cells with low number of genes that could represent artefacts. Cells with more than 10% of mitochondrial RNA in vitro and 0.25% in the graft data were removed to control for cell stress. The default method for identifying variable genes was used and the number of counts and mitochondrial genes was regressed out to remove biases due to these two variables. Data was normalized and transformed according to default methods in Scanpy. To reduce data dimensionality the 50 most significant PCs with 10 neighbors were used to construct a K-nearest neighbors (KNN) graph in vitro while in the graft dataset 20PCs and 20 neighbors were used. Batch effects of every sample were corrected using BBKNN^15^. The Leiden algorithm was used to define cell communities with a resolution of 1.0 in the in vitro data and 0.9 in the graft data. The uniform manifold approximation and projection (UMAP) was used to visualize the data in two dimensions. Cell communities were then annotated based on canonical markers. PAGA^16^ in the Scanpy package was used to measure the connectivity between the different cell types, all cell/types were considered except unknown cells. Only connections above the 0.2 thresholds were considered in the graph construction. For the *TH* or *NR4A2* filtered cells scran version 1.22.1 was used for data processing. The default 2000 highly variable genes were selected, and the first 50 principal components were extracted from the cell matrix. Subsequently, the shared nearest neighbors were calculated from the principal components using buildSNNGraph of R software scran. Clusters were identified using the walktrap algorithm, with the function cluster_walktrap of R implementation of the igraph package version 1.3.5. The uniform manifold approximation and projection (UMAP) was performed. Differential gene expression was performed via the Seurat package. Cluster pathway analysis was performed via clusterProfiler package version 4.2.2.

### Similarity score

To assess specific enrichments in boost and boost plus derived cells to authentic dopamine neurons of the midbrain we compared our datasets to human fetal (both in vitro and graft) and adult (only graft as A9 and A10 neurons are discernible in the graft) midbrain dopamine neurons single-cell data. We first calculated the top differentially expressed genes that define fetal dopamine neurons^12^ and adult^13^ A9 (considering all SOX6 DA neurons in the reference adult data) and A10 (considering all CALB1 positive DA neurons in the reference adult data) dopamine neurons using a Wilcoxon rank-sum test. P-values were then adjusted for multiple testing using the Benjamini-Hochberg test. These top 100 genes were used to compute a score of similarity for each cell in our dataset to this highly specific gene-list by considering the average expression of these genes in each cell in vitro or in the graft subtracted the average expression of a random set of genes in the dataset. We then plotted the score of each cell per protocol in boxplots and tested for differences in enrichment between boost and boost plus DA neurons by comparing distributions between protocols using the Mann-Whitney rank test and then adjusting p-values using the Benjamini-Hochberg test.

### Animals

All animal procedures and maintenance at Memorial Sloan Kettering Cancer Center (MSKCC) were approved by our Institutional Animal Care and Use Committee (IACUC) and following NIH guidelines. Athymic nude rats (NIH-Foxn1rnu, 6-8 weeks old, female, Charles Rivers Laboratory) and NSG mice (NOD.Cg-Prkdcscid Il2rgtm1Wjl/SzJ, 6-8 weeks old, male, Jackson Laboratory) were included in the studies.

### 6-OHDA lesioning and cell grafting

The animals were anesthetized by Isoflurane during the surgeries. For athymic nude rats, to establish unilateral medial forebrain bundle lesions of the nigro-striatal pathway, 6-OHDA solution (3.6 mg/ml in 0.2% ascorbic acid and 0.9% saline, Millipore) was stereotactically injected to (2.5μl, Tooth bar set at −2.4, AP −4.4 mm, ML −1.2, VL −7.8; 3 μl, Tooth bar set at +3.4, AP −4.0 mm, ML −0.8, VL −8.0). For the cell injections, human ESC-derived mDA progenitors (day 16) were resuspended at 100,000 ± 10,000 cells per microliter in transplantation medium consisting of neurobasal medium with 200 mM L-glutamine and 100 mM ascorbic acid (AA), 0.1% Kedbumin. The cell suspension were delivered to 4 deposits (1 µL per deposit) into the rat striatum (AP: +1mm, ML: −2.8, VL: −4.7, −4.6, −4.5 and −4.4 mm from dura) or 2 deposits along the DV axis (1 µL per deposit) into the mouse striatum ([AP] +0.5mm, [ML] +/-1.8mm, [DV] -3.4 to -3.3 mm from dura) at the rate of 0.5-1 μl/min via a motorized stereotaxic injector (Model 53311, Stoelting company, IL, USA). The syringe was kept in place for 5 minutes then slowly withdrawn at 1 mm/min.

Each surgery did not exceed more than 30 minutes per animal and surgery time for the entire cohort was within 8 - 10 hours post cell preparation. All cells used for transplantation studies underwent proper quality control (QC) metrics prior to injection such as immunofluorescence, intracellular flow, qPCR, and trypan blue or AOPI viability assays. All animal work was performed under the approval of the Institutional Animal Care and Use Committee (IACUC) at Memorial Sloan Kettering Cancer Center.

### Behavior tests

Amphetamine-induced rotation, ladder rung walking test, adhesive removal task were performed before transplantation, and at 1.5, 3, 4.5, 6 months after transplantation. The animals were habituated for 30 minutes before the behavior tests. For the amphetamine-induced rotation test, the rats were injected intraperitoneally with D-Amphetamine in saline (Sigma, 5mg/kg). The rotations were recorded for 40 minutes and automatically counted by Ethovision XT 16. The data were presented as (Ipsilateral-contralateral) rotations per minute. The ladder rung walking test was performed on the foot misplacement corridor (Panlab) with irregular rung arrangements (Metz et al., 2009). The percentage of missed steps out of the total number of steps was calculated. The adhesive removal task was performed with the adaptation that the adhesive tape was applied onto the forepaws (Fleming et al., 2013). The time the animal took to remove the tape off the left paw was recorded.

### Tissue Processing, Immunohistochemistry (IHC), image Processing and stereological analysis

The animals were perfused with 4% PFA. Brain tissues were dissected and post-fixed with 4% PFA for 12 hours, then changed to 30% sucrose in 0.01M PBS for 24 hours. The tissues were embedded in O.C.T (Sakura Finetek USA, Inc.) and cryosectioned at 30 μm. The primary antibodies included hNCAM (1:100), anti-TH (1:400), anti-hNA (1:100), anti-FOXA2 (1:100), Engrailed 1 (EN1, 1:25), dopamine transporter (DAT, 1:200), anti-GIRK2 (1:100), anti-ALDH1A1 (1:200), and anti-Calbindin (1:200). The cells or sections were washed with 0.1M PBS (Invitrogen) for 3 times of 5 min, then blocked with 1%BSA-0.3% Triton-PBS for 1hr. The primary antibodies were incubated overnight at 4°C. After washing with 0.01M PBS for 3 times of 5 min, the cells or tissue sections were incubated with appropriate fluorescent (Alexa 488, 568, 647) conjugated second antibodies (1:400, ThermoFisher Scientific) for 1 hour at room temperature. After rinsing with PBS, the nuclei were counterstain by DAPI. For DAB staining performed in MSKCC molecular cytology core facility, slides were loaded into Leica Bond RX and pretreated with EDTA-based epitope retrieval ER2 solution (Leica, AR9640) for 20 mins at 100°C. The sections were incubated with TH antibody (1ug/ml) at RT for 60 mins and subsequently with Leica Bond Polymer (anti-rabbit HRP) (included in Polymer Refine Detection Kit (Leica, DS9800)) for 8 min. Mixed DAB reagent (Polymer Refine Detection Kit) was applied for 10 mins. The sections were counterstained with Hematoxylin (Refine Detection Kit) for 10 mins. After staining, sample slides were washed in water, dehydrated using ethanol gradient (70%, 90%, 100%), washed three times in HistoClear II (National Diagnostics, HS-202), and mounted in Permount (Fisher Scientific, SP15).

The stained slides were scanned on a Panoramic Scanner (3DHistech, Budapest, Hungary) using a 20x/0.8NA objective in MSKCC molecular cytology core facility. The pictures were taken by Caseviewer 2.4 software (3DHistech Ltd, Budapest, Hungary). Stereological estimation by Stereo Investigator (MBF Bioscience, Vermont). The number of cells was determined using the optical fractionator probe while the volume of graft was analyzed using the Cavalieri estimation function.

### Statistical analysis

In all studies, animals were randomized into the different groups. Data were represented as mean ± SD unless indicated as mean± SEM. The number of cases in groups are specified in figure legends. One-way ANOVA was applied, and unpaired t test was used between two groups with equal SDs. Welch’s correction was applied to data with unequal SDs. Probability (p) values of less than 0.05 were considered statistically significant. All statistical analyses were analyzed by GraphPad Prism 9. To reduce the observation bias, the individuals who performed the experiments were blinded to group assignments in all procedures. Animals without any brain grafts in the cell group due to technical issues during transplantation were not included in the graft analyses.

| **Antibodies** |  |  |
| --- | --- | --- |
| Anti-hNA | Abcam | Cat# ab190710 |
| anti-tyrosine hydroxylase (Anti-TH) | Pel freeze | Cat# P40101-150, RRID:AB_2313713 |
| anti-TH | Pel freeze | Cat# P60101-150, RRID:AB_461070 |
| Anti-TH | Millipore | Cat#AB152, RRID: AB_390204 |
| Anti-FoxA2 | R&D system | Cat#AF2400, RRID:AB_2294104 |
| Anti-human specific Ki67 (anti-hKi67) | Agilent | Cat#GA62661-2, RRID:AB_2687921 |
| Anti-hNCAM (human specific) | Santa Cruz | Cat# SC106, RRID: AB_627128 |
| Anti-Engrailed 1 | Novus Biotechnology | Cat# NBP2-57757, RRID:AB_11015169 |
| Anti-DAT | Santa Cruz | Cat#sc-58517, RRID: AB_782391 |
| Sheep Anti-human Col1A1 | R&D | Cat#AF6220,RRID:AB_10891543 |
| Anti-GIRK2 | Alomone labs | Cat# APC_006-GPRRID: AB_2756604 |
| Anti-Calbindin | Cell signaling | Cat# 13176RRID: AB_2687400 |
| Anti-ALDH1A1 | Santa Cruz Biotechnology | Cat# sc-22589RRID: AB_2224026 |
| Anti-hKu80 (human specific) | Cell signaling Technology | Cat#2180sRRID: AB_2218736 |
| Anti-Prealbumin antibody | Abcam | Cat#ab92469, RRID:AB_1056381 |

## Data availability

All sc/snRNA-seq data have been deposited in the ArrayExpress database at EMBL-EBI (www.ebi.ac.uk/arrayexpress/) under accession no. E-MTAB-14729.

## Acknowledgments

We thank members of the Studer and Tabar lab for discussions on the manuscript. We would also like to thank the Flow Cytometry core, the Molecular Cytology core, the Integrated Genomic Operation (IGO) at MSKCC for outstanding technical support. Cores are supported by the NCI Cancer Center Support Grant (P30 CA08748), Cycle for Survival, and the Marie-Josée and Henry R. Kravis Center for Molecular Oncology. The work was supported in part by NIH grants 1R01 NS118067-01A1 (L.S., D.B.), R01NS126588 (V.T.), support by the National Research Foundation of Korea (NRF) graft funded by the Korea government (MSIT) (No. RS-2024-00351442) to T.W.K., support from the JPB foundation and from BlueRock Therapeutics to L.S.

## Author contributions

Conceptualization, T.W.K., J.P., V.T., and L.S. Writing – Original Draft, T.W.K., J.P., V.D.B., and L.S Dopamine neuron differentiation and characterization, T.W.K., S.Y.K., E.H., R.W. In vivo transplantation and analysis of the data, J.P., S.Y.K., L.R.P., S.J., Z.A.M., N.C., and S.A.D; Bioinformatics Analysis and interpretation of the data, V.D.B., F.C., D.Y.., H.S.C., and D.B; Electrophysiology and dopamine release experiments and analysis of the data, S.J.C., A.K.F., and E.V.M; Funding Acquisition, L.S., V.T., T.W.K., and D.B. All authors provided feedback on editing the manuscript. Co-first authorship was determined by the independent, and complementary leadership of the three co-first authors in hPSC differentiation study (T.W.K.), preclinical model development and analysis (J.P.), and computational analyses in vitro and in vivo (V.D.B.)

## SUPPLEMENTARY FIGURE LEGEND

**Fig. S1.**
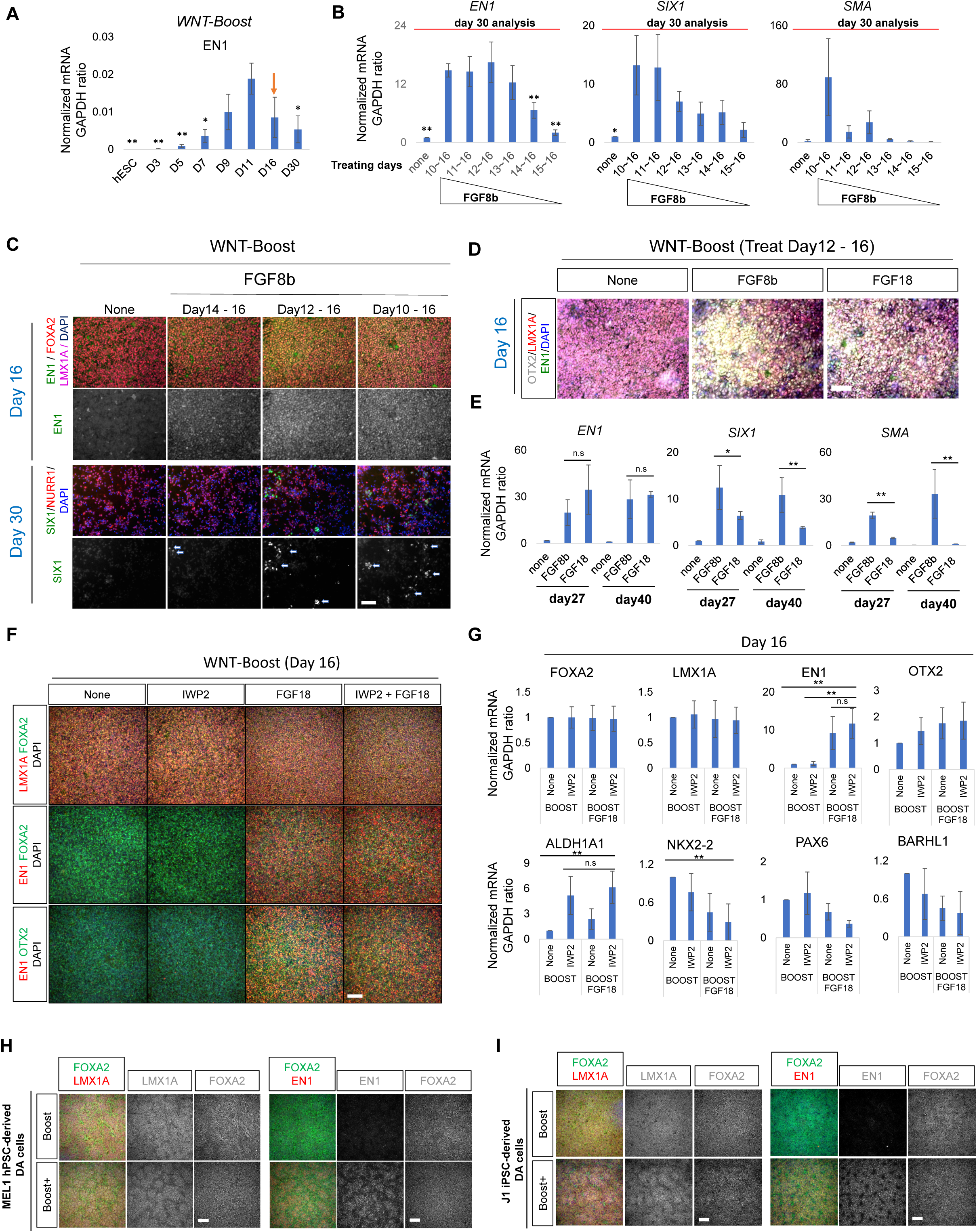
FGF18 and IWP2 treatment improves EN1+FOXA2+LMX1A+OTX2 mDA cells while reducing SIX1 and SMA1 cells across diverse hPSCs. **(A)** qRT-PCR analysis of mDA differentiated cells for *EN1* during differentiation from hPSC using the Boost protocol. **(B)** qRT-PCR analysis of mDA differentiated cells for mDA neuron marker, *EN1* and non-mDA contaminating markers, *SIX1* and *SMA1* at day30 with different duration treatment of FGF8b, from day10, 11, 12, 13, 14, and 15 to day16 on the Boost protocol. **(C)** Representative Immunofluorescent image of mDA differentiated cells for FOXA2 / LMX1A / EN1 and SIX1 / NURR1 at day16 and day30 with different duration treatment of FGF8b from day10, 12, and 14 to day16 on the Boost protocol. (**D**) Representative Immunofluorescent image of mDA differentiated cells for mDA markers, such as EN1, OTX2, and LMX1A at day16 with treatment of FGF18 from day12 to day16 on the Boost protocol. **(E)** qRT-PCR assay of mDA differentiated cells at day27 and day40 with treatment of FGF8b and FGF18 from day12 to day16 on the Boost protocol. **(F)** Representative Immunofluorescent image of mDA differentiated cells for mDA markers at day16 with treatment of IWP2 only, FGF18 only, and IWP2+FGF18 from day12 to day16 on the Boost protocol. **(G)** qRT-PCR analysis of mDA differentiated cells at day16 from (**F**). **(H, I)** Representative Immunofluorescent image of mDA differentiated cells for mDA markers at day16 derived from MEL1 hESC (**H**) and J1 iPSC (**I**) by the Boost and Boost+ protocol. Data are represented as mean±SD. *P<0.05, ** P<0.001. p-value was calculated *via* comparison from Day11 in **A**, and from 12∼16 in **B**.

**Fig S2.**
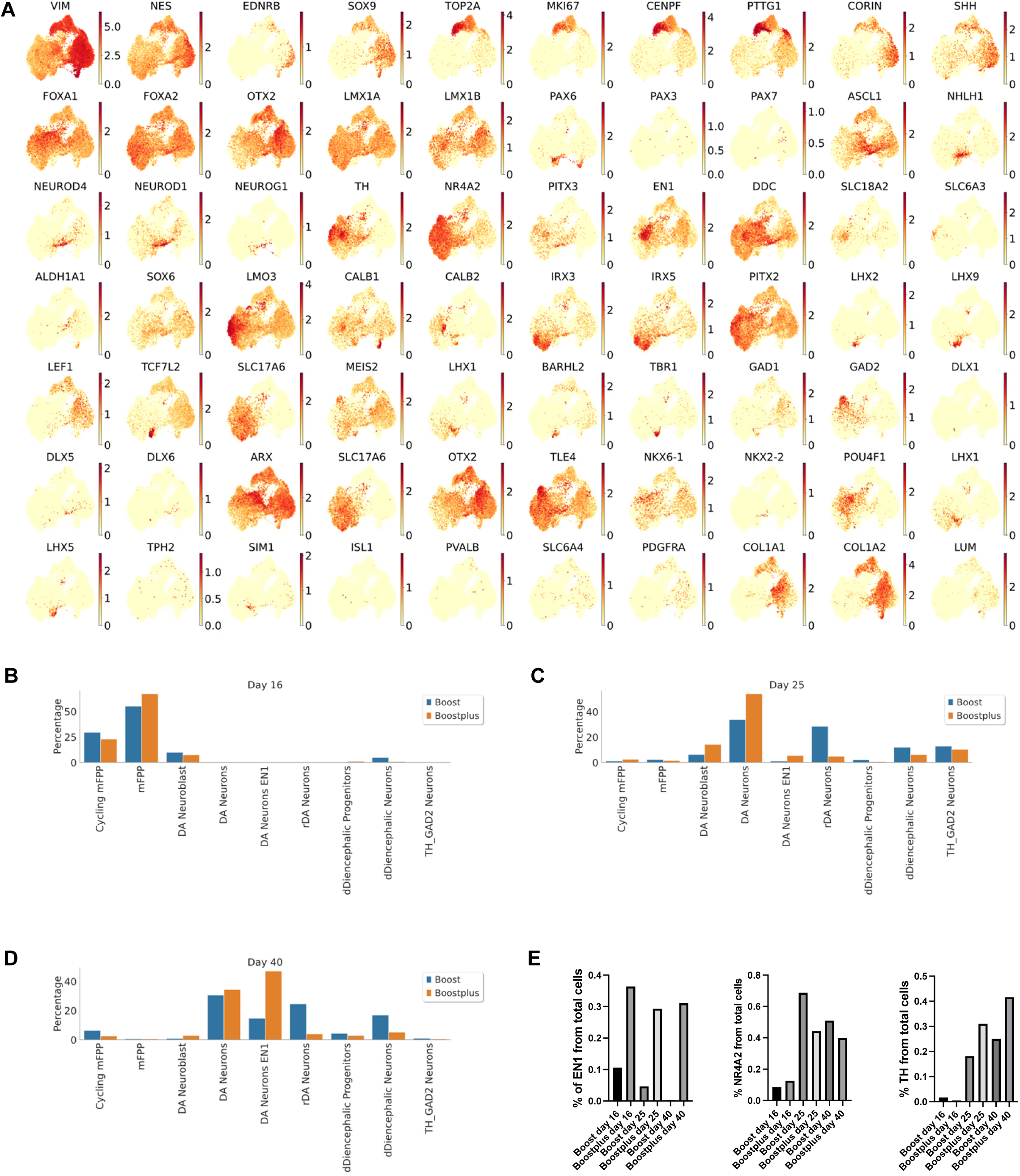
scRNA-seq analysis during mDA differentiation combined at day16, 25, 40 in Boost and Boost+ conditions *in vitro*. **(A)** UMAP plot showing expression levels of markers of general neuronal progenitors (*VIM*,*NES*,*EDNRB*, *SOX9*), cycling progenitors (*TOP2A*, *MKI67*, *CENPF*, *PTTG1*), midbrain floor progenitors (*CORIN*, *SHH*, *FOXA1*, *FOXA2*, *OTX2*, *LMX1A*, *LMX1B*), dorsal diencephalic progenitors (*PAX6*, *PAX3*, *PAX7*), neuroblasts (*ASCL1*, *NHLH1*, *NEUROD4*, *NEUROD1*, *NEUROG1*), general dopamine neurons (*TH*, *NR4A2*, *PITX3*, *EN1*, *DDC*, *SLC18A2*, *SLC6A3*), A9 mDA neurons (*ALDH1A1*, *SOX6*, *LMO3*), A10 mDA neurons (*CALB1*, *CALB2*), subthalamic neurons (*IRX3*, *IRX5*, *PITX2*), thalamic neurons (*LHX2*, *LHX9*, *LEF1*, *TCF7L2*, *SLC17A6*), pretectal neurons (*MEIS2*, *LHX1*, *BARHL2*, *TBR1*), vLGN interneurons (*GAD1*, *GAD2*, *DLX1*, *DLX5*, *DLX6*, *ARX*, *SLC17A6*, *OTX2*, *TLE4*), red nucleus (*NKX6-1*, *NKX2-2*, *POU4F1*, *LHX1*, *LHX5*, *TPH*), OMTN (*SIM1*, *ISL1*, *PVALB*), serotonergic neurons (*SLC6A4*) and VLMCs (*PDGFRA*, *COL1A1*, *COL1A2*, *LUM*). **(B-D)** Percentage of cell types in each protocol at day 16 **(B)**, day 25 **(C)** and day 40 **(D)**. **(E)** Percentage of cells expressing *EN1*, *NR4A2* and *TH* from the scRNA-seq

**Fig S3.**
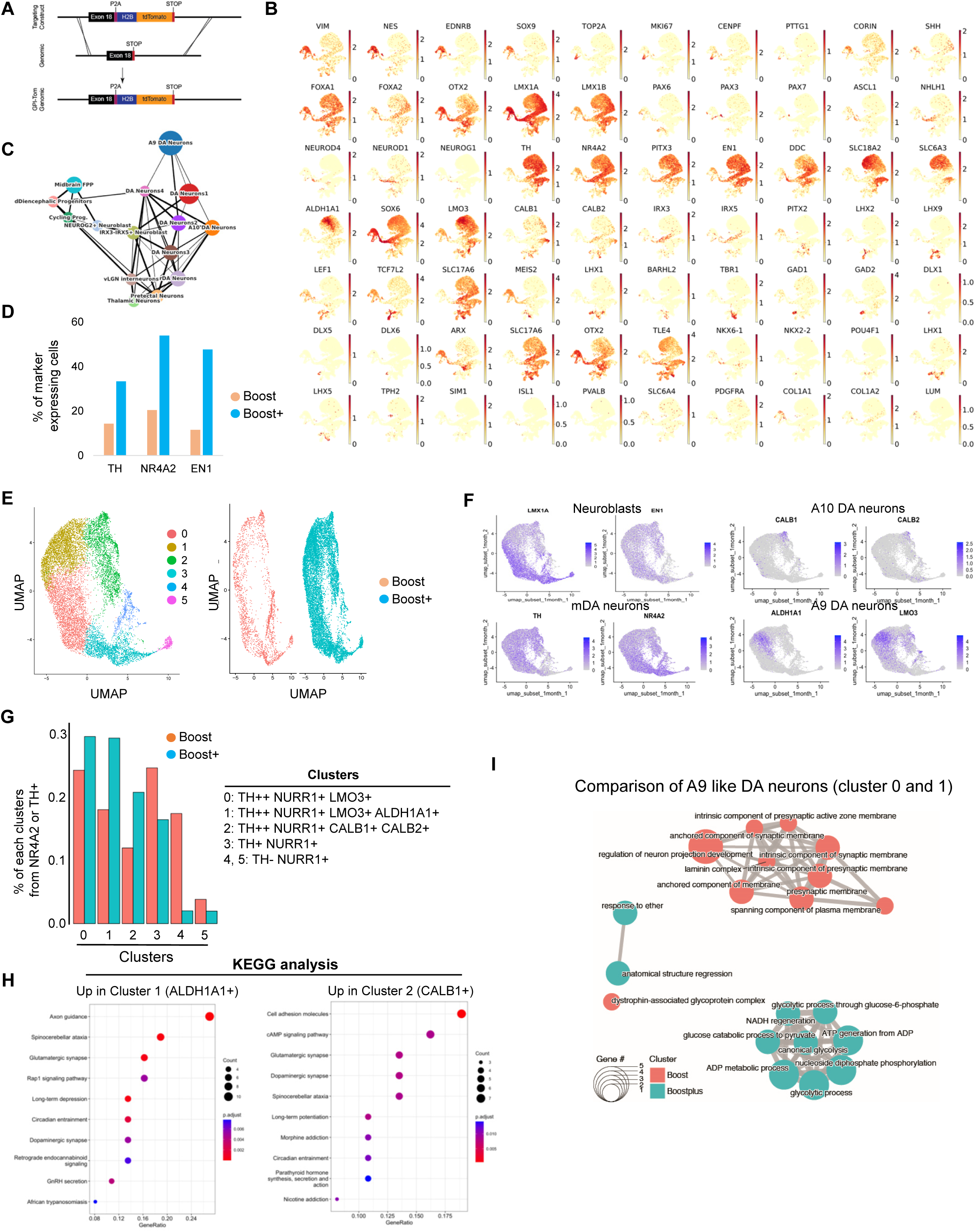
Characterization of cell type composition within grafts using snRNA-seq. **(A)** Schematic illustration of generation reporter line to isolate human grafted mDA cells expressing Tomato for snRNA-seq. **(B)** UMAP plot showing expression levels of markers of general neuronal progenitors (*VIM*,*NES*,*EDNRB*, *SOX9*), cycling progenitors (*TOP2A*, *MKI67*, *CENPF*, *PTTG1*), midbrain floor progenitors (*CORIN*, *SHH*, *FOXA1*, *FOXA2*, *OTX2*, *LMX1A*, *LMX1B*), dorsal diencephalic progenitors (*PAX6*, *PAX3*, *PAX7*), neuroblasts (*ASCL1*, *NHLH1*, *NEUROD4*, *NEUROD1*, *NEUROG1*), general dopamine neurons (*TH*, *NR4A2*, *PITX3*, *EN1*, *DDC*, *SLC18A2*, *SLC6A3*), A9 mDA neurons (*ALDH1A1*, *SOX6*, *LMO3*), A10 mDA neurons (*CALB1*, *CALB2*), subthalamic neurons (*IRX3*, *IRX5*, *PITX2*), thalamic neurons (*LHX2*, *LHX9*, *LEF1*, *TCF7L2*, *SLC17A6*), pretectal neurons (*MEIS2*, *LHX1*, *BARHL2*, *TBR1*), vLGN interneurons (*GAD1*, *GAD2*, *DLX1*, *DLX5*, *DLX6*, *ARX*, *SLC17A6*, *OTX2*, *TLE4*), red nucleus (*NKX6-1*, *NKX2-2*, *POU4F1*, *LHX1*, *LHX5*, *TPH*), OMTN (*SIM1*, *ISL1*, *PVALB*), serotonergic neurons (*SLC6A4*) and VLMCs (*PDGFRA*, *COL1A1*, *COL1A2*, *LUM*). **(C)** Abstracted graph showing the relationships between cell states. **(D-I)** Characterization of *TH*+ and *NR4A2*+ cells from snRNA-seq in the graft. Percent of cells expressing gene markers TH, NR4A2, and EN1 separated by the Boost and Boost+ **(D)**. UMAP visualization of clusters of combined Boost and Boost+ cells (**E, left**) and UMAP visualization of clusters separated by Boost and Boost+ cells (**E, right**). UMAP visualization of selected gene markers highly correlated with cell types neuroblast, mDA neurons, A9 mDA neurons, and A10 mDA neurons **(F).** Percentage of each cluster of the *NR4A2* or *TH*+ filter by the Boost and Boost+ **(G).** KEGG-GO analysis of upregulated pathway in high expression of ALDH1A1 and CALB1 clusters **(H).** Plot showing pathways (or biological processes) enriched in the engrafted A9 mDA cells generated using the Boost (red) or Boost+ (green) protocol **(I)**.

**Fig. S4.**
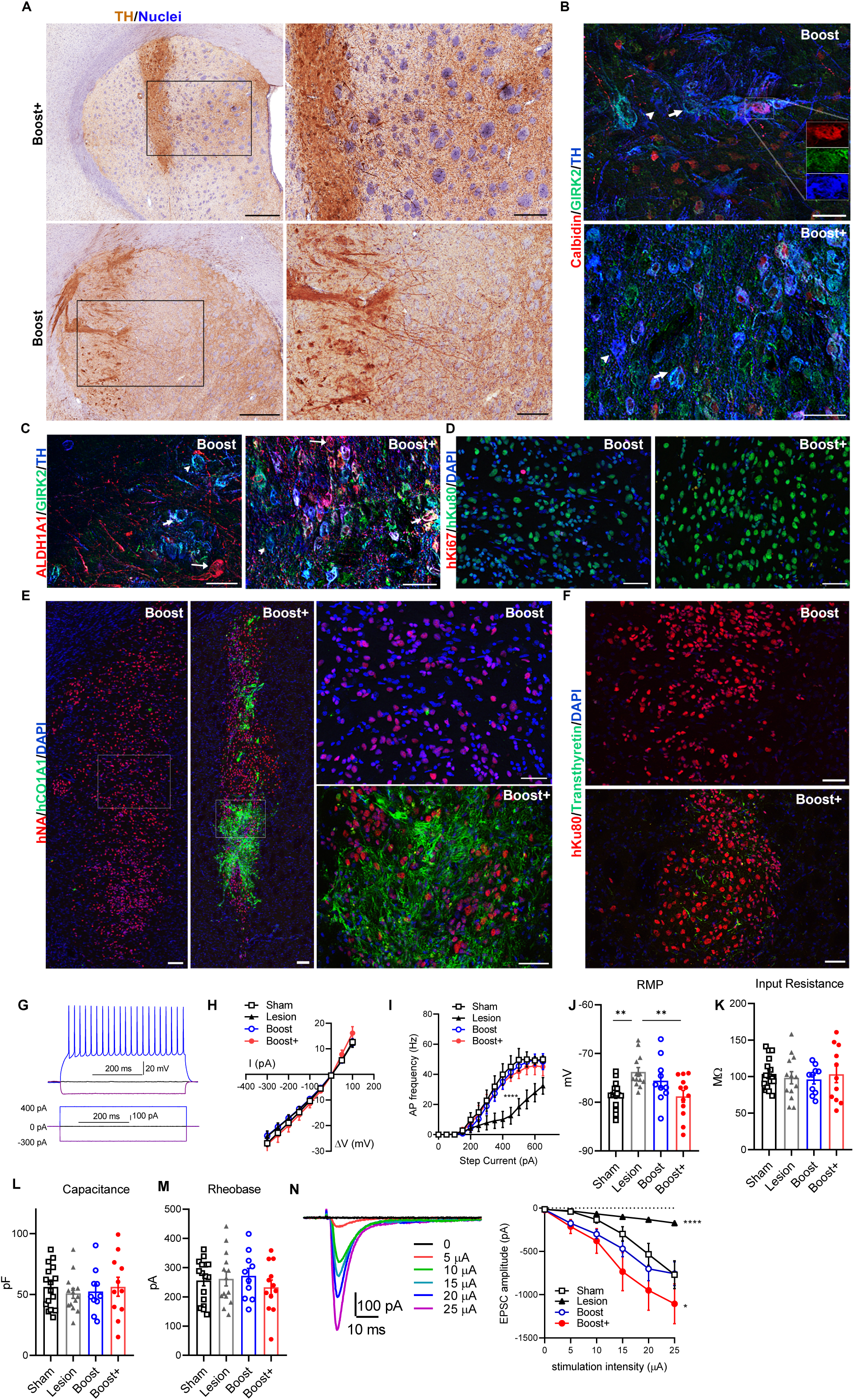
Characterization of Boost and Boost+ grafts in unilateral 6-OHDA lesioned rat striatum. **(A)** IHC for TH shows graft derived dopaminergic processes in the host striatum in Boost and Boost+ cell grafts. Scale bar represents 500 µm on the left panels while 200 µm on the right panels. Nuclei were counterstained with hematoxylin. **(B)** Representative images of IHC for TH+ cells co-labeling GIRK2 and CALBINDIN in Boost and Boost+ grafts. Arrowheads point to TH+ cells that do not express GIRK2 and CALBIDIN while arrows point to TH+ cells that express GIRK2. In boost grafts, the TH ^+^ cell that expresses both CALBIDIN and GIRK2 is outlined in white lines and shown as split channels in an inset. Note that some GIRK2+TH+ cells express CALBINDIN. **(C)** Representative images of IHC for TH ^+^ cells co-labeling GIRK2 and ALDH1A1 in Boost and Boost+ grafts. Arrowheads point to TH ^+^ GRIK2 ^+^ cells that do not express ALDH1A1 while short arrows point to GIRK2+ TH+ cells that express ALDH1A1. Long arow points to ALDH1A1 ^+^ TH ^+^ cells that do not express GIRK2. **(D)** Representative images of immunohistochemistry (IHC) for human specific Ki67 (hKi67) and human specific marker hKu80 in Boost and Boost+ grafts. The nuclei were counterstained with DAPI. Scale bar=50 µm. **(E)** Representative images of IHC for human specific COL1A1 (hCOL1A1) and human specific marker hKu80 in Boost and Boost+ grafts. The nuclei were counterstained with DAPI. Scale bar represents 100 µm on the left panels while 50 µm on the right panels. **(F)** Representative images of IHC for transthyretin (Prealbumin) and human specific hKu80 in Boost and Boost+ grafts. The nuclei were counterstained with DAPI. Scale bar=50 µm. **(G-N)** Whole-cell recordings from striatal spiny projection neurons (SPN) proximal to the mDA neurons graft. Examples of SPN response to current injections **(G)**, voltage-current curve **(H)** and dependence of action potential frequency on the amplitude of injected current **(I).** ****- p<0.0001 from other groups by 2-way ANOVA. Membrane properties of SPNs, including resting membrane potential (J, n=10-16 cells), input resistance **(K)**, membrane capacitance **(L)** and rheobase **(M)**. **(N)** Representative EPSC traces evoked by electrical stimulation of the corpus callosum in the presence of GABAA antagonist picrotoxin (**left**) and dependence of EPSC amplitude on stimulation current intensity (**right**). **** and *- different from all other groups by 2-way ANOVA with p<0.0001 and 0.05, correspondingly.

**Fig. S5.**
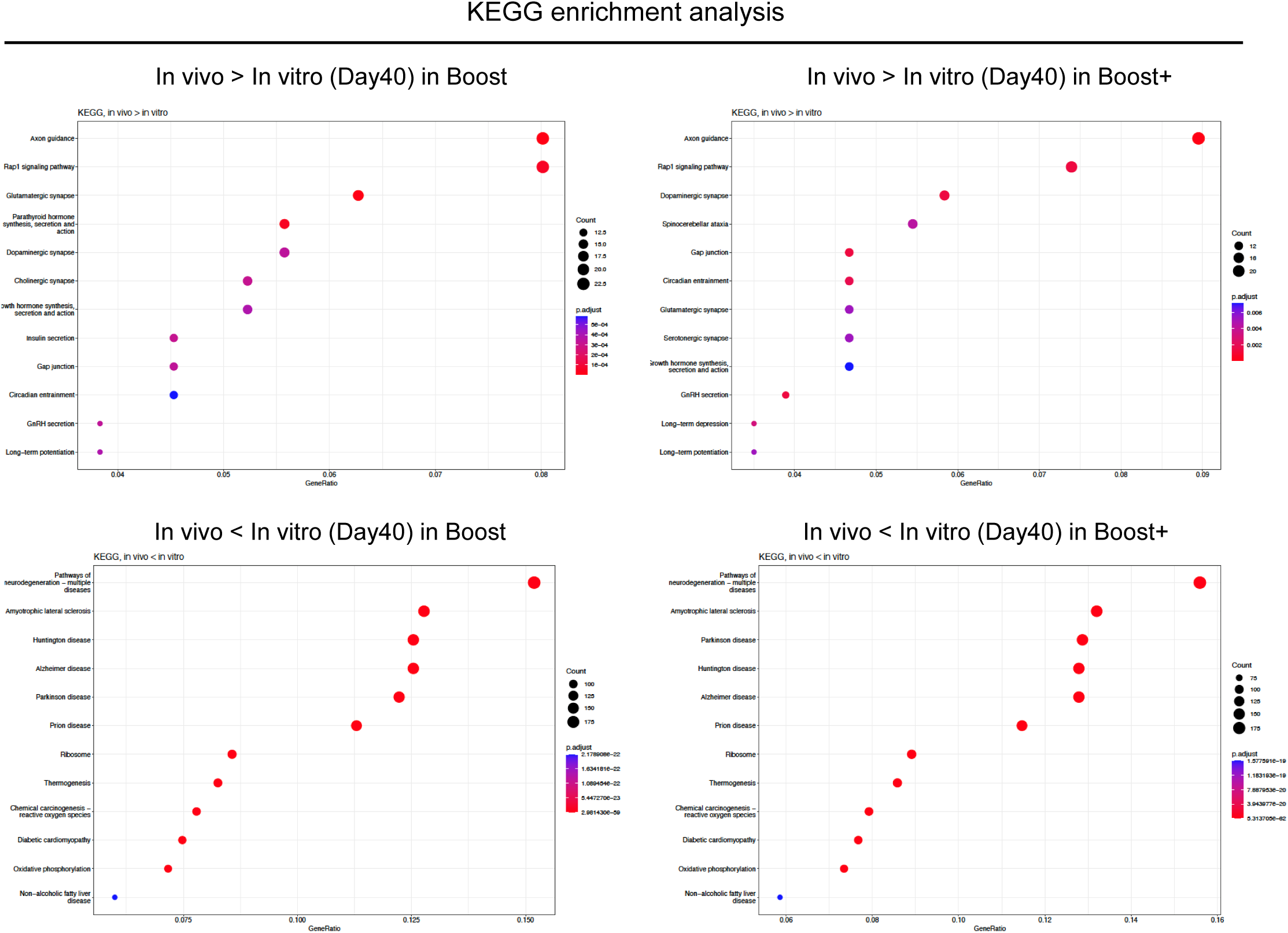
KEGG-GO analysis of NURR1+ or TH+ mDA neurons comparing *in vivo* grafts (snRNA-seq) and age-compatible *in vitro* mDA neurons (scRNAseq, Day40) in the Boost and Boost+ conditions.

## Uncategorized References

1. Poewe W, Seppi K, Tanner CM, Halliday GM, Brundin P, Volkmann J, et al. Parkinson disease. Nat Rev Dis Primers. 2017;3:17013.

2. Bloem BR, Okun MS, and Klein C. Parkinson’s disease. Lancet. 2021;397(10291):2284–303.

3. Barker RA, Drouin-Ouellet J, and Parmar M. Cell-based therapies for Parkinson disease-past insights and future potential. Nat Rev Neurol. 2015;11(9):492–503.

4. Tabar V, and Studer L. Pluripotent stem cells in regenerative medicine: challenges and recent progress. Nat Rev Genet. 2014;15(2):82–92.

5. Lindvall O, and Bjorklund A. Cell therapy in Parkinson’s disease. NeuroRx. 2004;1(4):382–93.

6. Barker RA, and consortium T. Designing stem-cell-based dopamine cell replacement trials for Parkinson’s disease. Nat Med. 2019;25(7):1045–53.

7. Kim TW, Koo SY, and Studer L. Pluripotent Stem Cell Therapies for Parkinson Disease: Present Challenges and Future Opportunities. Front Cell Dev Biol. 2020;8:729.

8. Parmar M, Grealish S, and Henchcliffe C. The future of stem cell therapies for Parkinson disease. Nat Rev Neurosci. 2020;21(2):103–15.

9. Kim TW, Piao J, Koo SY, Kriks S, Chung SY, Betel D, et al. Biphasic Activation of WNT Signaling Facilitates the Derivation of Midbrain Dopamine Neurons from hESCs for Translational Use. Cell Stem Cell. 2021;28(2):343–55 e5.

10. Song B, Cha Y, Ko S, Jeon J, Lee N, Seo H, et al. Human autologous iPSC-derived dopaminergic progenitors restore motor function in Parkinson’s disease models. J Clin Invest. 2020;130(2):904–20.

11. Doi D, Magotani H, Kikuchi T, Ikeda M, Hiramatsu S, Yoshida K, et al. Pre-clinical study of induced pluripotent stem cell-derived dopaminergic progenitor cells for Parkinson’s disease. Nat Commun. 2020;11(1):3369.

12. Kirkeby A, Nolbrant S, Tiklova K, Heuer A, Kee N, Cardoso T, et al. Predictive Markers Guide Differentiation to Improve Graft Outcome in Clinical Translation of hESC-Based Therapy for Parkinson’s Disease. Cell Stem Cell. 2017;20(1):135–48.

13. Kikuchi T, Morizane A, Doi D, Magotani H, Onoe H, Hayashi T, et al. Human iPS cell-derived dopaminergic neurons function in a primate Parkinson’s disease model. Nature. 2017;548(7669):592-6.

14. Piao J, Zabierowski S, Dubose BN, Hill EJ, Navare M, Claros N, et al. Preclinical Efficacy and Safety of a Human Embryonic Stem Cell-Derived Midbrain Dopamine Progenitor Product, MSK-DA01. Cell Stem Cell. 2021;28(2):217–29 e7.

15. Schweitzer JS, Song B, Herrington TM, Park TY, Lee N, Ko S, et al. Personalized iPSC-Derived Dopamine Progenitor Cells for Parkinson’s Disease. N Engl J Med. 2020;382(20):1926–32.

16. Kirkeby A, Nelander J, Hoban DB, Rogelius N, Bjartmarz H, Novo Nordisk Cell Therapy R, et al. Preclinical quality, safety, and efficacy of a human embryonic stem cell-derived product for the treatment of Parkinson’s disease, STEM-PD. Cell Stem Cell. 2023;30(10):1299–314 e9.

17. Release BTP. BlueRock’s Phase I study with bemdaneprocel in patients with Parkinson’s disease meets primary endpoint. https://www.prnewswire.com/news-releases/bluerocks-phase-i-study-with-bemdaneprocel-in-patients-with-parkinsons-disease-meets-primary-endpoint-301910863.html.

18. Takahashi J. iPS cell-based therapy for Parkinson’s disease: A Kyoto trial. Regen Ther. 2020;13:18–22.

19. Gantner CW, de Luzy IR, Kauhausen JA, Moriarty N, Niclis JC, Bye CR, et al. Viral Delivery of GDNF Promotes Functional Integration of Human Stem Cell Grafts in Parkinson’s Disease. Cell Stem Cell. 2020;26(4):511–26 e5.

20. Kriks S, Shim JW, Piao J, Ganat YM, Wakeman DR, Xie Z, et al. Dopamine neurons derived from human ES cells efficiently engraft in animal models of Parkinson’s disease. Nature. 2011;480(7378):547-51.

21. Kirkeby A, Grealish S, Wolf DA, Nelander J, Wood J, Lundblad M, et al. Generation of regionally specified neural progenitors and functional neurons from human embryonic stem cells under defined conditions. Cell Rep. 2012;1(6):703–14.

22. Cha Y, Park TY, Leblanc P, and Kim KS. Current Status and Future Perspectives on Stem Cell-Based Therapies for Parkinson’s Disease. J Mov Disord. 2023;16(1):22–41.

23. Kee N, Volakakis N, Kirkeby A, Dahl L, Storvall H, Nolbrant S, et al. Single-Cell Analysis Reveals a Close Relationship between Differentiating Dopamine and Subthalamic Nucleus Neuronal Lineages. Cell Stem Cell. 2017;20(1):29–40.

24. Nouri N, and Awatramani R. A novel floor plate boundary defined by adjacent En1 and Dbx1 microdomains distinguishes midbrain dopamine and hypothalamic neurons. Development. 2017;144(5):916–27.

25. Tiklova K, Nolbrant S, Fiorenzano A, Bjorklund AK, Sharma Y, Heuer A, et al. Single cell transcriptomics identifies stem cell-derived graft composition in a model of Parkinson’s disease. Nat Commun. 2020;11(1):2434.

26. Xu P, He H, Gao Q, Zhou Y, Wu Z, Zhang X, et al. Human midbrain dopaminergic neuronal differentiation markers predict cell therapy outcomes in a Parkinson’s disease model. J Clin Invest. 2022;132(14).

27. Li H, Jiang H, Li H, Li L, Yan Z, and Feng J. Generation of human A9 dopaminergic pacemakers from induced pluripotent stem cells. Mol Psychiatry. 2022;27(11):4407–18.

28. Nishimura K, Yang S, Lee KW, Asgrimsdottir ES, Nikouei K, Paslawski W, et al. Single-cell transcriptomics reveals correct developmental dynamics and high-quality midbrain cell types by improved hESC differentiation. Stem Cell Reports. 2023;18(1):337–53.

29. Oosterveen T, Garcao P, Moles-Garcia E, Soleilhavoup C, Travaglio M, Sheraz S, et al. Pluripotent stem cell derived dopaminergic subpopulations model the selective neuron degeneration in Parkinson’s disease. Stem Cell Reports. 2021;16(11):2718–35.

30. Maxwell SL, Ho HY, Kuehner E, Zhao S, and Li M. Pitx3 regulates tyrosine hydroxylase expression in the substantia nigra and identifies a subgroup of mesencephalic dopaminergic progenitor neurons during mouse development. Dev Biol. 2005;282(2):467–79.

31. Hwang DY, Ardayfio P, Kang UJ, Semina EV, and Kim KS. Selective loss of dopaminergic neurons in the substantia nigra of Pitx3-deficient aphakia mice. Brain Res Mol Brain Res. 2003;114(2):123–31.

32. La Manno G, Gyllborg D, Codeluppi S, Nishimura K, Salto C, Zeisel A, et al. Molecular Diversity of Midbrain Development in Mouse, Human, and Stem Cells. Cell. 2016;167(2):566–80 e19.

33. Liu G, Yu J, Ding J, Xie C, Sun L, Rudenko I, et al. Aldehyde dehydrogenase 1 defines and protects a nigrostriatal dopaminergic neuron subpopulation. J Clin Invest. 2014;124(7):3032–46.

34. Wu J, Kung J, Dong J, Chang L, Xie C, Habib A, et al. Distinct Connectivity and Functionality of Aldehyde Dehydrogenase 1a1-Positive Nigrostriatal Dopaminergic Neurons in Motor Learning. Cell Rep. 2019;28(5):1167–81 e7.

35. Tiklova K, Bjorklund AK, Lahti L, Fiorenzano A, Nolbrant S, Gillberg L, et al. Single-cell RNA sequencing reveals midbrain dopamine neuron diversity emerging during mouse brain development. Nat Commun. 2019;10(1):581.

36. Poulin JF, Zou J, Drouin-Ouellet J, Kim KY, Cicchetti F, and Awatramani RB. Defining midbrain dopaminergic neuron diversity by single-cell gene expression profiling. Cell Rep. 2014;9(3):930–43.

37. Cuevas-Diaz Duran R, Gonzalez-Orozco JC, Velasco I, and Wu JQ. Single-cell and single-nuclei RNA sequencing as powerful tools to decipher cellular heterogeneity and dysregulation in neurodegenerative diseases. Front Cell Dev Biol. 2022;10:884748.

38. Alvarez-Fischer D, Fuchs J, Castagner F, Stettler O, Massiani-Beaudoin O, Moya KL, et al. Engrailed protects mouse midbrain dopaminergic neurons against mitochondrial complex I insults. Nat Neurosci. 2011;14(10):1260–6.

39. Nordstroma U, Beauvais G, Ghosh A, Pulikkaparambil Sasidharan BC, Lundblad M, Fuchs J, et al. Progressive nigrostriatal terminal dysfunction and degeneration in the engrailed1 heterozygous mouse model of Parkinson’s disease. Neurobiol Dis. 2015;73:70–82.

40. Sgado P, Alberi L, Gherbassi D, Galasso SL, Ramakers GM, Alavian KN, et al. Slow progressive degeneration of nigral dopaminergic neurons in postnatal Engrailed mutant mice. Proc Natl Acad Sci U S A. 2006;103(41):15242–7.

41. Lahti L, Peltopuro P, Piepponen TP, and Partanen J. Cell-autonomous FGF signaling regulates anteroposterior patterning and neuronal differentiation in the mesodiencephalic dopaminergic progenitor domain. Development. 2012;139(5):894–905.

42. Itoh N, and Ornitz DM. Fibroblast growth factors: from molecular evolution to roles in development, metabolism and disease. J Biochem. 2011;149(2):121–30.

43. Liu A, Li JY, Bromleigh C, Lao Z, Niswander LA, and Joyner AL. FGF17b and FGF18 have different midbrain regulatory properties from FGF8b or activated FGF receptors. Development. 2003;130(25):6175–85.

44. Arenas E. Wnt signaling in midbrain dopaminergic neuron development and regenerative medicine for Parkinson’s disease. J Mol Cell Biol. 2014;6(1):42–53.

45. Chilov D, Sinjushina N, Saarimaki-Vire J, Taketo MM, and Partanen J. beta-Catenin regulates intercellular signalling networks and cell-type specific transcription in the developing mouse midbrain-rhombomere 1 region. PLoS One. 2010;5(6):e10881.

46. Tang M, Villaescusa JC, Luo SX, Guitarte C, Lei S, Miyamoto Y, et al. Interactions of Wnt/beta-catenin signaling and sonic hedgehog regulate the neurogenesis of ventral midbrain dopamine neurons. J Neurosci. 2010;30(27):9280–91.

47. Lipiec MA, Bem J, Kozinski K, Chakraborty C, Urban-Ciecko J, Zajkowski T, et al. TCF7L2 regulates postmitotic differentiation programmes and excitability patterns in the thalamus. Development. 2020;147(16).

48. Brozko N, Baggio S, Lipiec MA, Jankowska M, Szewczyk LM, Gabriel MO, et al. Genoarchitecture of the Early Postmitotic Pretectum and the Role of Wnt Signaling in Shaping Pretectal Neurochemical Anatomy in Zebrafish. Front Neuroanat. 2022;16:838567.

49. Wisniewska MB, Misztal K, Michowski W, Szczot M, Purta E, Lesniak W, et al. LEF1/beta-catenin complex regulates transcription of the Cav3.1 calcium channel gene (Cacna1g) in thalamic neurons of the adult brain. J Neurosci. 2010;30(14):4957–69.

50. Ferran JL, Sanchez-Arrones L, Sandoval JE, and Puelles L. A model of early molecular regionalization in the chicken embryonic pretectum. J Comp Neurol. 2007;505(4):379–403.

51. Mastick GS, Davis NM, Andrew GL, and Easter SS, Jr. Pax-6 functions in boundary formation and axon guidance in the embryonic mouse forebrain. Development. 1997;124(10):1985–97.

52. Ding Q, Balasubramanian R, Zheng D, Liang G, and Gan L. Barhl2 Determines the Early Patterning of the Diencephalon by Regulating Shh. Mol Neurobiol. 2017;54(6):4414–20.

53. Siletti K, Hodge R, Mossi Albiach A, Lee KW, Ding SL, Hu L, et al. Transcriptomic diversity of cell types across the adult human brain. Science. 2023;382(6667):eadd7046.

54. Bhaduri A, Andrews MG, Mancia Leon W, Jung D, Shin D, Allen D, et al. Cell stress in cortical organoids impairs molecular subtype specification. Nature. 2020;578(7793):142-8.

55. Satija R, Farrell JA, Gennert D, Schier AF, and Regev A. Spatial reconstruction of single-cell gene expression data. Nat Biotechnol. 2015;33(5):495–502.

56. Golding B, Pouchelon G, Bellone C, Murthy S, Di Nardo AA, Govindan S, et al. Retinal Input Directs the Recruitment of Inhibitory Interneurons into Thalamic Visual Circuits. Neuron. 2014;81(6):1443.

57. Sabbagh U, Govindaiah G, Somaiya RD, Ha RV, Wei JC, Guido W, et al. Diverse GABAergic neurons organize into subtype-specific sublaminae in the ventral lateral geniculate nucleus. J Neurochem. 2021;159(3):479–97.

58. Niehaus JL, Cruz-Bermudez ND, and Kauer JA. Plasticity of addiction: a mesolimbic dopamine short-circuit? Am J Addict. 2009;18(4):259–71.

59. Kamath T, Abdulraouf A, Burris S, Gazestani V, Nadaf N, Vanderburg C, et al. A molecular census of midbrain dopaminergic neurons in Parkinson’s disease. bioRxiv. 2021:2021.06.16.448661.

60. Park S, Park CW, Eom JH, Jo MY, Hur HJ, Choi SK, et al. Preclinical and dose-ranging assessment of hESC-derived dopaminergic progenitors for a clinical trial on Parkinson’s disease. Cell Stem Cell. 2023.

61. Doi D, Samata B, Katsukawa M, Kikuchi T, Morizane A, Ono Y, et al. Isolation of human induced pluripotent stem cell-derived dopaminergic progenitors by cell sorting for successful transplantation. Stem Cell Reports. 2014;2(3):337–50.

62. Lehnen D, Barral S, Cardoso T, Grealish S, Heuer A, Smiyakin A, et al. IAP-Based Cell Sorting Results in Homogeneous Transplantable Dopaminergic Precursor Cells Derived from Human Pluripotent Stem Cells. Stem Cell Reports. 2017;9(4):1207–20.

63. Samata B, Doi D, Nishimura K, Kikuchi T, Watanabe A, Sakamoto Y, et al. Purification of functional human ES and iPSC-derived midbrain dopaminergic progenitors using LRTM1. Nat Commun. 2016;7:13097.

64. Metz GA, and Whishaw IQ. The ladder rung walking task: a scoring system and its practical application. J Vis Exp. 2009(28).

65. Metz GA, and Whishaw IQ. Drug-induced rotation intensity in unilateral dopamine-depleted rats is not correlated with end point or qualitative measures of forelimb or hindlimb motor performance. Neuroscience. 2002;111(2):325–36.

66. Kim TW, Koo SY, Riessland M, Cho H, Chaudhry F, Kolisnyk B, et al. TNF-NFkB-p53 axis restricts in vivo survival of hPSC-derived dopamine neuron. bioRxiv. 2023.

67. Miller JD, Ganat Y, Kishinevsky S, Bowman R, Liu B, Tu E, et al. Human iPSC-based Modeling of Late-Onset Disease via Progerin-induced Aging. Cell Stem Cell. 2013;(in press).

68. Krishnaswami SR, Grindberg RV, Novotny M, Venepally P, Lacar B, Bhutani K, et al. Using single nuclei for RNA-seq to capture the transcriptome of postmortem neurons. Nat Protoc. 2016;11(3):499–524.

69. Braun E, Danan-Gotthold M, Borm LE, Lee KW, Vinsland E, Lonnerberg P, et al. Comprehensive cell atlas of the first-trimester developing human brain. Science. 2023;382(6667):eadf1226.

